# RegVar: Tissue-specific Prioritization of Noncoding Regulatory Variants

**DOI:** 10.1101/2021.04.17.440295

**Authors:** Hao Lu, Luyu Ma, Cheng Quan, Lei Li, Yiming Lu, Gangqiao Zhou, Chenggang Zhang

**Affiliations:** Beijing Institute of Radiation Medicine, State Key Laboratory of Proteomics, Beijing 100850, China

**Keywords:** Variant prioritization, Expression regulation, Deep neural network

## Abstract

Noncoding genomic variants constitute the majority of trait-associated genome variations; however, identification of functional noncoding variants is still a challenge in human genetics, and a method systematically assessing the impact of regulatory variants on gene expression and linking them to potential target genes is still lacking. Here we introduce a deep neural network (DNN)-based computational framework, RegVar, that can accurately predict the tissue-specific impact of noncoding regulatory variants on target genes. We show that, by robustly learning the genomic characteristics of massive variant-gene expression associations in a variety of human tissues, RegVar vastly surpasses all current noncoding variants prioritization methods in predicting regulatory variants under different circumstances. The unique features of RegVar make it an excellent framework for assessing the regulatory impact of any variant on its putative target genes in a variety of tissues. RegVar is available as a webserver at http://regvar.cbportal.org/.

## Introduction

Trait-associated genetic variants usually lie in noncoding genomic regions [1, 2], and interpretation of functional noncoding variants is crucial for revealing the underlying genetic architecture and molecular mechanism of complex traits and diseases. Several methods have been developed to discriminate pathogenic variants from non-pathogenic ones using genomic sequences, functional annotations and evolutionary features, such as CADD [3], GWAVA [4], DeepSEA [5], LINSIGHT [6], etc. A common feature of these methods is that they focus on identifying rare pathogenic variants, which were thought to have stronger impact on human traits and diseases than common variants [7]. However, emerging evidences suggest that the majority of heritability for complex traits is likely to be explained by a substantial number of common regulatory variants with small additive effect sizes, in combination with a relatively smaller contribution from rare variants of moderate effect sizes [8–10]. Thus, a model that can distinguish both common and rare regulatory variants will provide new perspectives on the regulatory basis of complex traits.

Current pathogenic variant prioritization models are not suitable for identifying regulatory variants. A recent survey of existing methods for prioritizing noncoding variants showed that, although they achieved high precision in identifying pathogenic variants under certain circumstances, their performance in identifying regulatory variants was very poor [11]. This is because prioritization of regulatory variants is an even greater challenge than that of pathogenic ones. First, regulatory variants generally have weaker impact on gene expression compared to pathogenic ones, so it is more difficult to discriminate them from background, especially from adjacent non-functional variants sharing similar epigenetic marks. Second, it is challenging to link regulatory variants to their target genes, which can be located far away from its regulator. Third, it is a challenge to establish tissue or cell type-specific models that can predict the regulatory impact of variants under different biological conditions. A number of methods have been proposed to predict the effects of regulatory variants in recent years [12–14], which, however, have their limitations in their application. For example, ExPecto relies on epigenetic marks at gene promoters to monitor the regulatory impact of variants on gene expression and thus could only assess promoter-proximal variants [12]; TIVAN connects various genomic features to expression quantitative trait loci (eQTLs) to estimate a variant’s regulatory probability, but it was trained with promoter-proximal variants, which may introduce potential biases when applied to genome-wide variants prioritization [13]. Considering the vast majority of regulatory variants located far from the transcription start sites (TSSs) of target genes [15], a method that can robustly predict genome-wide regulatory variants as well as their potential target genes remains an urgent need.

Here we introduce a deep neural network (DNN)-based approach, RegVar, for the genome-wide assessment of the regulatory impact of noncoding variants on gene expression. RegVar has several key features: (i) it can predict both common and rare regulatory variants by learning their genomic characteristics from massive variant-gene associations in an unbiased manner; (ii) it predicts not only regulatory variants but their target genes by jointly learning the genomic patterns of both variants and genes and the chromatin interactions between them; (iii) it predicts the tissue-specific effects of variants by training models in multiple tissues with respective genomic patterns; and (iv) it can achieve excellent prediction accuracy by utilizing large training sets and deep learning algorithm. We show that RegVar outperforms existing prioritization methods in identifying regulatory variants and noncoding pathogenic variants from different backgrounds in various tissues. RegVar is available as a webserver at http://regvar.cbportal.org/.

## Materials and methods

### Datasets

To construct the positive datasets, significant eVariant-eGene associations in 17 human tissues, which were also incorporated in the Roadmap Epigenomics projects [2], were obtained from GTEx V7 release [16] (**Figure 1** and Table S1). Single-nucleotide variants (SNVs)-gene associations were selected and further filtered by removing eVariants not marked by DNase I hypersensitive sites (DHSs) annotations, which was demonstrated to be a key epigenetic marker of causal variants [16]. Associations in sex chromosomes were also removed. For tissues of which the numbers of significant associations exceed 100,000 (esophagus mucosa, lung, skeletal muscle, and whole blood), we randomly selected 100,000 associations, as we found a larger size cannot improve model performance (Figure S1). The final number of positive associations for each tissue was shown in Table S1. For negative datasets, four datasets were constructed, including: (i) *random-variant* set of shuffled SNV-gene pairs where eVariants were replaced by random SNVs located <=1 Mb from the eGene TSS; (ii) *mirrored-variant* set of shuffled pairs where eVariants were replaced by random SNVs located at similar distance (error <= 1kb) but the opposite side of the eGene TSS; (iii) *neighboring-variant* set of shuffled pairs where eVariants were replaced by random SNVs located adjacent (<= 1 kb) to the positive ones; (iv) *random-gene* set of shuffled SNV-gene pairs where eGenes were replaced by gene TSSs located <= 1 Mb of the eVariants. We selected a maximum distance at 1 Mb between SNV and TSS in the datasets (i) and (iv), for it was observed that all positive SNV-TSS pairs had a distance less than 1 Mb (Figure S2). To determine the ratio between positive and negative datasets, we assessed different ratios, including 1:1, 1:2, 1:3, 1:5, and 1:10, and found there was no significant difference of performances among five models (Figure S3). Thus, we selected a ratio of 1:1 between the positive and negative datasets to efficiently train the models. Variants in negative datasets were selected from the dbSNP build 146 data after removing the shared variants between GTEx and dbSNP datasets. Since eQTL variants are biased toward high frequency variants (Figure S4), to ensure that our results were not influenced by the differences in minor allele frequency (MAF) between the positive variants and negative controls, we defined additional sets of MAF-matched negative controls for GTEx liver dataset, by the same strategy as the first three control datasets (i-iii) described above.

**Figure 1.**
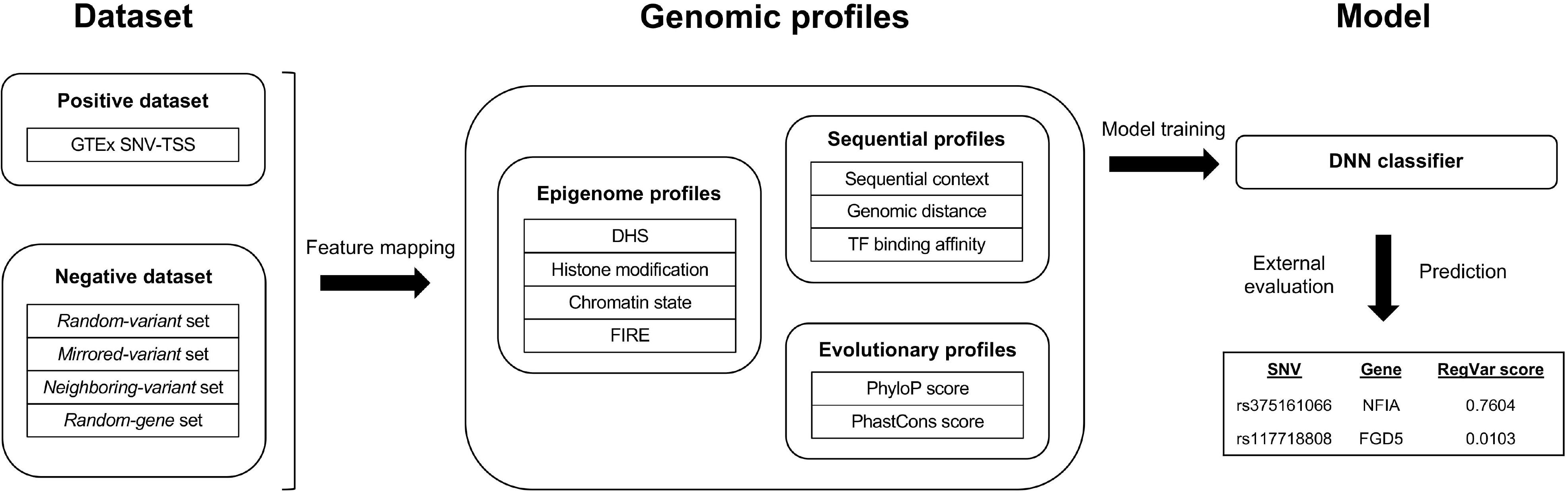
A flowchart showing the workflow of RegVar.

### Annotation profiles

We used three major categories of genomic profiles, including sequential, epigenetic and evolutionary profiles (Table S2), to annotate our datasets using a customized pipeline.

#### Sequential profiles

Sequential profiles consisted of 2-mer prefix and postfix and local 5-mer GC content of SNV and TSS, SNV-caused transcription factor binding site (TFBS) affinity changes, genomic distance between SNV and TSS and the orientations of SNV and TSS.

To calculate TFBS affinity changes caused by variants, we obtained the position frequency matrices of 602 TFs from the TRANSFAC [17] (523 TFs) and JASPAR [18] (79 TFs) databases. TFMscan [19] was used to locate putative TFBS motifs by scanning genomic DNA both forward and backward using these position frequency matrices. A stringent threshold of *P*-value < 4.5E-5 was used to determine significant motifs. Variants located within these motifs were determined using BEDTools [20]. The TFBS affinity were calculated as described [21]. Specifically, the corrected probabilities of observing a given nucleotide in a specific locus were calculated as follows:

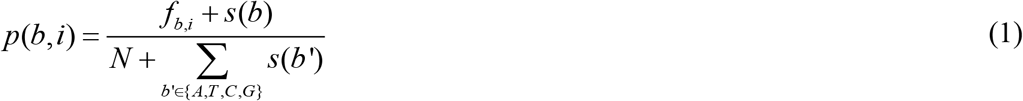

Where *b* represents one specific base among A, T, C, and G, *i* is the index of the site, *f_b,i_* is the counts of base *b* in site *i*, *N* is the sum of counts of four bases, and *s(b)* is the pseudocount function. Here we assumed *s(b)* to be 1/4 for each of the four bases, then

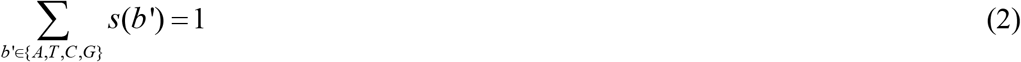

Hence, the corresponding PWM can be constructed as:

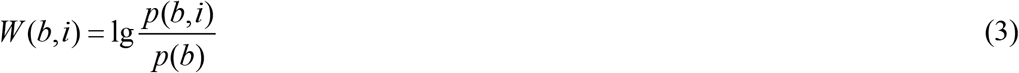

where *p(b)* is the background probability of base *b* (assumed to be 1/4 for four bases). The TFBS affinity is calculated with

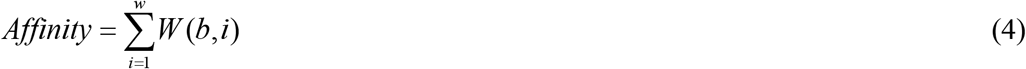

where *w* is the width of a PWM. We then calculated the average affinity change between reference and alteration alleles as follows:

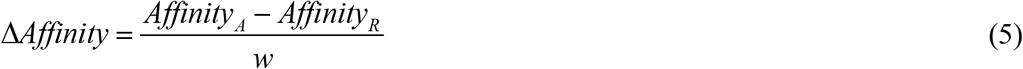

where *Affinity_R_* and *Affinity_A_* are evaluated binding affinity with the reference and alteration alleles, respectively. Variants located within two or more TFBS motifs were assigned with the *ΔAffinity* score with the maximum absolute value among all *ΔAffinity* scores of the affected motifs and variants not located at any TFBS motif were assigned a *ΔAffinity* score of 0.

#### Epigenetic profiles

Epigenetic profiles consisted of 31 histone modifications from the Roadmap Epigenomics project [2], 25 chromatin states produced by ChromHMM [22], and frequently interacted regions (FIREs) annotations from Hi-C study [23].

#### Evolutionary profiles

Evolutionary profiles consisted of vertebrate, placental mammal and primate phyloP [24] and phastCons [25] scores based on the 46-way whole-genome alignment, and vertebrate phyloP and phastCons scores based on the 100-way whole-genome alignment.

All annotations were expressed in genomic coordinates for the GRCh37/hg19 assembly of the human genome. Boolean variables were used to indicate if SNV or TSS overlapped with chromatin marks (1) or not (0). For categorical annotations, all n-level categorical values were first encoded to binary values and then converted to several individual Boolean flags. For continuous annotations, feature values were scaled to the range of [0, 1]. More exactly, distance to TSS was scaled by

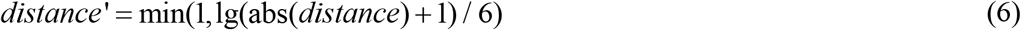

phyloP scores scaled by

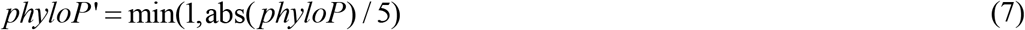

and *ΔAffinity* scores scaled by

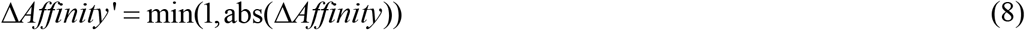

### Model design and training

We built a DNN-based classifier to model our dataset. The basic model in RegVar is a fully connected neural network, in which each neuron in a layer receives inputs from all outputs of the previous layer, except that the first layer receives inputs from the original data matrix. Each layer in the network executes a linear transformation of the corresponding inputs to integrate information from the previous layer, followed by a non-linear transformation (namely the activation function) to rectify the linear result. Here we employed three fully connected layers with 500, 200, and 60 units respectively, and the most used rectified linear unit function (ReLU) as the activation function. Exactly, one fully connected layer computes

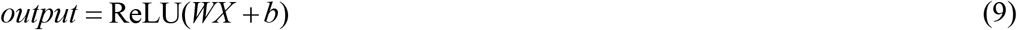

where *X* is the input, *W* is the weight matrix, *b* is the bias, and ReLU represents rectified linear function

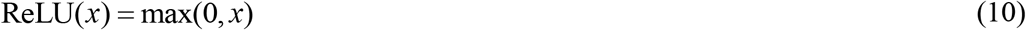

The layer following the third fully connected layer is the final output layer to make predictions about being a regulatory or non-regulatory variant on the specific gene, with scaled probability ranging from 0 to 1 using

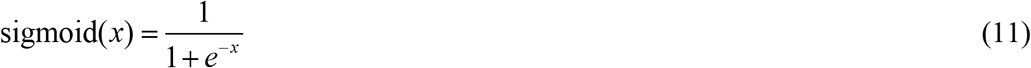

To train the model, we selected the cross entropy loss function as the objective function, which is defined as follows:

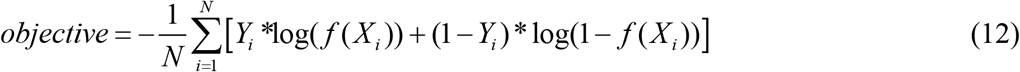

where *N* is the number of samples in the training set, and *i* is the index of each sample. *Y_i_* and *X_i_* represent the 0/1 label and the input features for sample *i*, respectively; and *f(X_i_)* represents the predicted probability output from the DNN model.

We conducted optimal search of hyper-parameters including the learning rate and dropout proportion. Learning rates were set at 0.001, 0.005, and 0.01; dropout proportions were set at 0, 0.3, and 0.5. We selected the combinations of learning rates and dropout proportions that achieved the highest prediction AUC in each of the four models (Table S3-S6).

All training programs were written in Python language, using a deep neural network implementation from the TensorFlow library.

### Model comparison

We used the average receiver operating characteristic (ROC) curves computed from 10-fold cross-validation to evaluate model performances. Specifically, each dataset comprising of the positive set and its negative counterpart was randomly split into a training set and a testing set in a 9:1 ratio; the RegVar model was trained on the training set and evaluated on the testing set. This process was repeated 10 times for each dataset, with independent sample split procedure each time.

Predictions of CADD (v1.3) [3], GWAVA (v1.0) [4], DeepSEA [5], LINSIGHT [6], ExPecto [12], and TIVAN [13], together with two ensemble methods, IW-scoring [26] and regBase [27] were used for model performance comparison in liver, hippocampus and whole blood datasets. The *random-variant*, *mirrored-variant* and *neighboring-variant* datasets were used for the evaluation, and *random-gene* dataset was excluded as the existing methods didn’t give prediction on potentially affected genes. In addition, ExPecto was excluded from the *random-variant* and *random-gene* models evaluation, because it focused on promoter-proximal variants thus resulted in too few samples for the evaluation. For CADD, DeepSEA, and IW-scoring, we ran the analysis using the corresponding online web services; for GWAVA, LINSIGHT, ExPecto, TIVAN, and regBase, we downloaded the precomputed scores from the corresponding source websites.

### Model external evaluation

We downloaded liver eQTLs from the exSNP website [28], hippocampus eQTLs from Schulz [29] and Ramasamy [30] eQTLs studies, and blood eQTLs from Westra [31] eQTLs meta-analysis to evaluate performances of trained models. We identified all SNV-TSS pairs and removed those overlapping with liver, hippocampus and whole blood eQTLs in GTEx dataset. For negative controls, all SNVs in the external positive datasets were removed from dbSNP build 146 and then four negative datasets were constructed, as described above in model training, for each of the three independent positive datasets. Also, the negative samples overlapping with the control sets used in model training were further removed to avoid any valid set contamination. Then we annotated each sample set with classifiers trained on GTEx eQTLs in the corresponding tissue and compared classification results with ROC curves for the first three sets with existing methods mentioned above.

Besides, we evaluated prediction capabilities of different methods on the liver and blood eQTLs data from Brown eQTLs analysis [32], which have been used to test the performance of TIVAN and regBase. We downloaded the compiled positive and negative sets for Brown eQTLs data from the regBase website and compared performance of different methods on datasets in liver and blood (all testing datasets are summarized in Table S7 and see Supplementary Methods for more details about data processing).

### RegVar score distribution

For each tissue, we trained an integrated RegVar model by pooling four negative datasets to take all conditions together. 17 integrated RegVar models were applied to annotate all possible SNV-gene pairs in chromosome 22. For each SNV in chromosome 22, we obtained TSSs of all genes located within 1 Mb of the variant locus and combined the variant with each of these TSSs as a possible eQTL pair. After mapped with all kinds of features, 65,844,726 sample pairs of 1,039,985 different SNVs were left and annotated with integrated RegVar models in 17 tissue types. For each variant, the maximum annotated score of all its possible eQTL pairs was set as the final RegVar score of the variant in each tissue. We next explored distribution patterns of RegVar scores of all these variants.

### Tissue-shared/tissue-specific regulatory variants

Stratified random sampling was performed to select 100,000 SNVs from 22 autosomes, and TSSs located within 1 Mb of each variant locus were identified and combined with the variant as a possible eQTL pair. After mapped with corresponding features, 3,703,900 sample pairs were left. RegVar scores were obtained in 17 tissue types and then converted to percentiles based on the corresponding merged training sets to make results comparable across different tissues. For a particular variant, the sample pair with the maximum percentile among all its possible eQTL pairs and across 17 integrated models was set to be the final sample pair. We obtained 17 tissue-specific percentiles of all final sample pairs to form a percentile matrix. Then K-means clustering, implemented by *kmeans* function in R language, was applied on the matrix to get tissue-specific and tissue-shared regulatory variant clusters. Four tissue-specific epigenetic features, namely DHS, H3K4me1, H3K4me3, and H3K27ac, were used to annotate these tissue-specific and tissue-shared regulatory variants.

## Results

### Prioritization of regulatory variants in 17 human tissues

To explore the influence of DHS filter on RegVar’s prediction capability, we first compared performance of models built from positive datasets without any filter, with the DHS filter, with the ATAC-seq filter, and both positive and negative datasets with the DHS filter in the liver dataset (Supplementary Methods), and found that models built from the positive dataset with the DHS filter showed the most robust performance in discriminating regulatory variants form different backgrounds (Figure S5). We then utilized the DHS-filter-based RegVar to predict the tissue-specific effects of genomic variants on gene expression in 17 human tissues. The averaged ROC curves across 17 tissues showed that RegVar predicted regulatory variants and their target genes with averaged AUCs of 0.965, 0.917, 0.693, and 0.929 for the four training datasets, respectively (**Figure 2** and Table S1). This result demonstrated RegVar could reliably discriminate positive regulatory variants from different negative backgrounds. We then evaluated the performances of existing methods CADD [3], GWAVA [4], DeepSEA [5], LINSIGHT [6], ExPecto [12], TIVAN [13], IW-scoring [26], and regBase [27] on the same tasks. For CADD, we used C-scores. For GWAVA, we used pathogenic scores with the corresponding control standards (namely, *unmatched*, *TSS*, and *region*). For DeepSEA, we used eQTL-probability scores. For IW-scoring, we used integrative scores without fitCons. For regBase, we used regBase_Common prediction scores. For the three tested tissues: liver, hippocampus, and whole blood, we found that only GWAVA, LINSIGHT, and IW-scoring could make valid predictions with AUCs of 0.668-0.764 for *random-variant* and 0.573-0.677 for *mirrored-variant* datasets, yet still much lower than that of RegVar (0.957-0.969 for *random-variant* and 0.884-0.945 for *mirrored-variant* sets), while other five methods failed to show significant power in distinguishing regulatory variants (**Figure 3**, Figure S6 and S7). For the *neighboring-variant* set, which is more challenging, none of the existing methods made valid predictions, compared to an AUC of 0.694-0.700 for RegVar. We additionally evaluate the prediction results of different methods by precision-recall curves (PRC), and still RegVar showed the superior performance against other methods (Figure S8, S9, and S10). After controlled for MAF, RegVar showed comparable prediction capabilities in discriminating eQTLs from MAF-matched benign variants, as demonstrated in liver samples (Figure S11), although with a slight decrease in the *neighboring-variant* set, and the other eight methods still showed low prediction capabilities as before.

**Figure 2.**
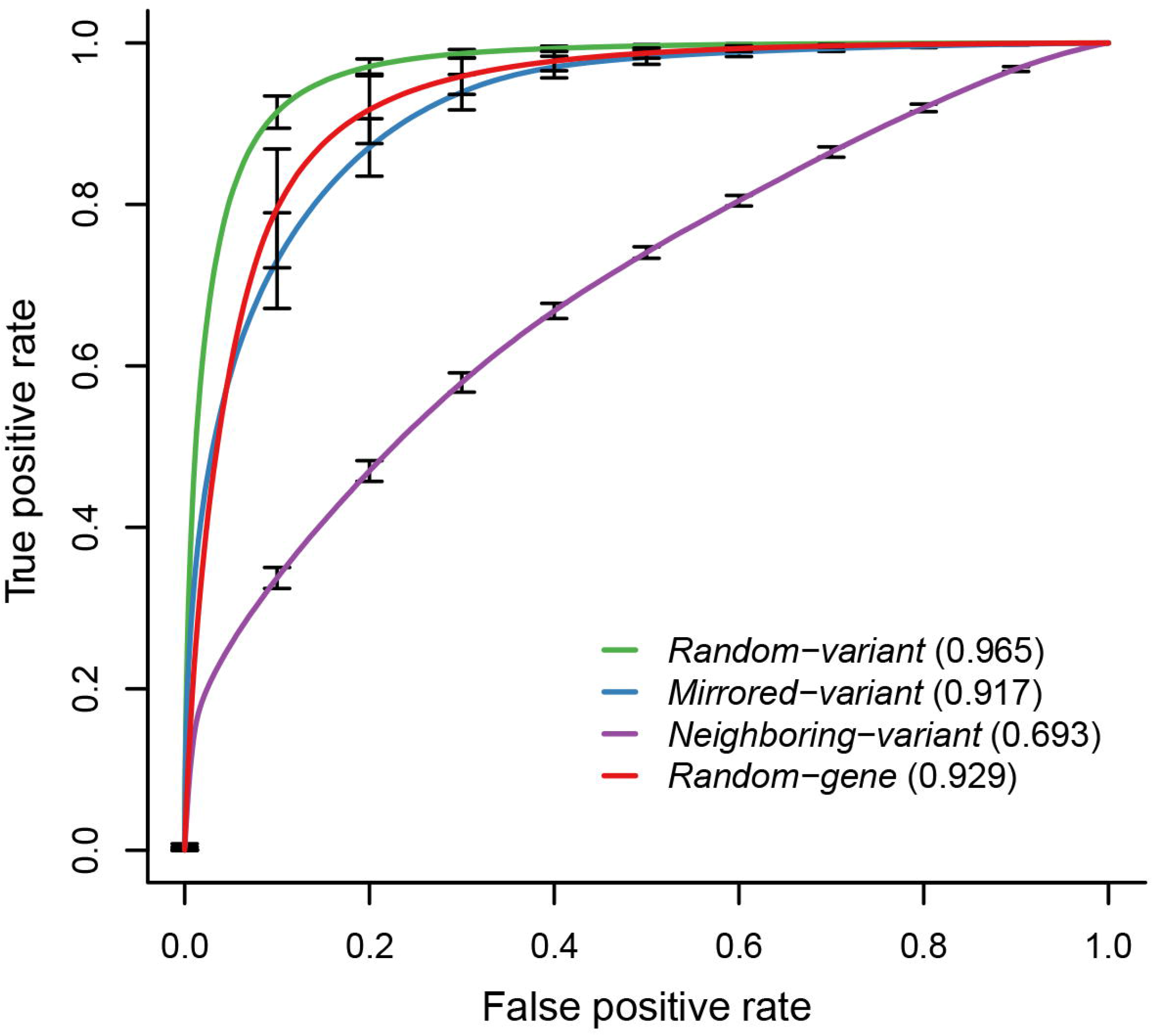
Average ROC curves for 10-fold cross-validation experiments of RegVar. For each of the four training sets, the ROC curves are averaged across 17 human tissues. Error bars represent the standard deviation averaged over tissues.

**Figure 3.**
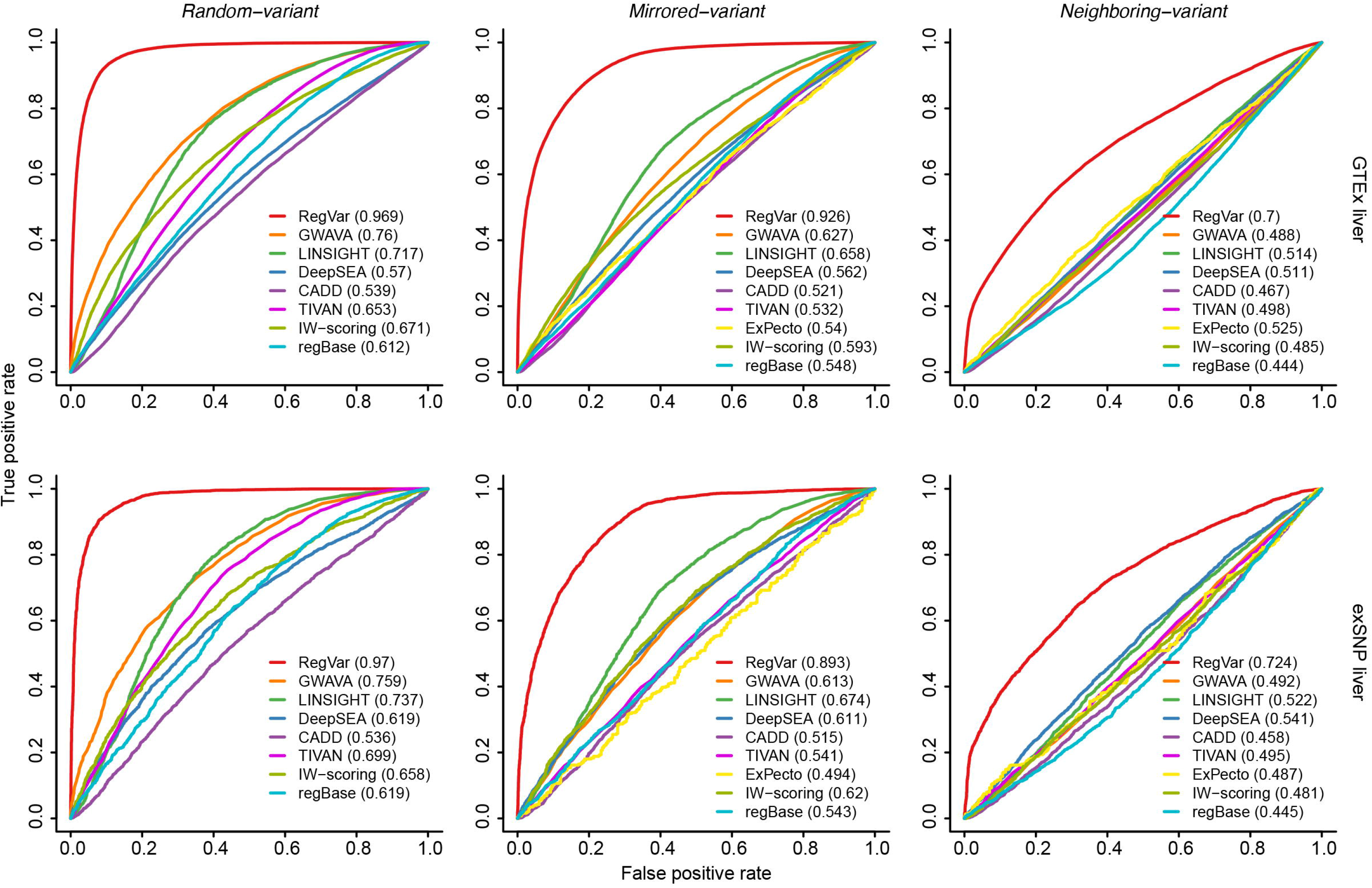
ROC curves of nine computational methods distinguishing regulatory variants from different backgrounds in liver. Results are shown for ROC curves from 10-fold cross-validation experiments in the GTEx liver eQTL dataset (top) and from external evaluation experiments in the exSNP liver eQTL dataset (bottom). Negative datasets were from either random selected variants (*random-variant* sets) (left), matched variants by distance but at the opposite side of the eGene TSS (*mirrored-variant* sets) (middle), or neighboring variants located adjacent (<= 1 kb) to the positive ones (*neighboring-variant* sets) (right). Negative datasets from random selected TSSs (*random-gene* sets) are not shown since other existing methods didn’t give prediction on potentially affected genes. Any overlap between the exSNP liver eQTLs and GTEx eQTLs and overlap between their corresponding negative sets were removed. ExPecto results are not shown for *random-variant* sets because it resulted in too few samples for ROC curve analysis.

To further confirm the results, we curated another three publicly available eQTL datasets of liver, hippocampus, and whole blood, from the exSNP website [28], Schulz [29] and Ramasamy [30] eQTL studies, and Westra [31] eQTL meta-analysis, respectively, as independent testing sets. We found RegVar models trained with GTEx datasets achieved almost equally accurate predictions in the three independent testing sets, while all other methods still didn’t show any obvious predictive powers in the independent datasets (Figure 3, Figure S6-S10). To assess the robustness of RegVar on imbalanced datasets, we then constructed independent validation sets for liver eQTLs form the exSNP database with the sample ratio at 1:1, 1:2, 1:3, 1:5, and 1:10. We found that RegVar trained on GTEx datatsets at the 1:1 ratio showed robust performance on both balanced and imbalanced datasets (Figure S12). In addition, we also evaluated performance of different methods on the Brown eQTL data in liver and blood, which have been used as testing data for regBase and TIVAN. Results showed that in both tissues, RegVar trained on GTEx datasets showed comparable performance (AUC = 0.858 and 0.901 for liver and blood sets, respectively) with regBase (AUC = 0.883 and 0.89 for liver and blood sets, respectively, equal to the AUCs reported in the regBase paper. We also showed that RegVar models trained on the Brown eQTL data achieved even higher AUCs on both datasets (AUC = 0.952 and 0.945 for liver and blood sets, respectively) compared with other methods (Figure S13 and S14).

To investigate the robustness of RegVar on different settings of negative data sampling in external evaluation, we constructed negative datasets for the exSNP testing set by randomly selecting variants at wider genome regions, including: (i) *random-variant* set comprising of random SNVs located <= 2 and 5 Mb from the eGene TSS; (ii) *mirrored-variant* set comprising of random SNVs with a distance error <= 2 and 5 kb; (iii) *neighboring-variant* set comprising of random SNVs located <= 2 and 5 kb to the positive ones; (iv) *random-gene* set comprising of random gene TSSs located <= 2 and 5 Mb of the eVariants. RegVar models trained before exhibited equal, or even slightly increased, prediction power in these independent negative controls selected from wider genome regions, whereas other methods again showed vary limited prediction performances (Figure S15). Altogether, these results demonstrated the outstanding performance of RegVar on predicting regulatory impact of noncoding variants.

We examined the feature importance of the four different models with Gini impurity in liver (Supplementary Methods). For *random-variant* and *mirrored-variant* models, the epigenetic patterns of variants were the most important feature sets, while for the *random-gene* model, the epigenetic and sequential profiles of TSS were the most important feature sets besides the distance between variant and TSS. Notably, for the *neighboring-variant* model, evolutionary and sequential profiles of variants became the most important feature sets (Figure S16). This is expected, as these features could provide information of single-base resolution, which is crucial for distinguishing regulatory variants from adjacent non-functional ones.

### RegVar score distribution

We further compared performances of different models trained on the GTEx liver eQTLs on each of negative datasets constructed for the independent exSNP testing set. Results showed that AUCs of models from other types of negative data decreased by 0.031-0.133, 0.041-0.091, 0.062-0.105 for *random-variant*, *mirrored-variant*, and *neighboring-variant* datasets, respectively, and that the integrated model obtained superior capability in all testing datasets besides the models trained on the same type of negative data (Figure S17). We then applied the integrated model to all SNVs in chromosome 22 (*n* = 1,039,985) in 17 types of tissues and measured their regulatory potentials with the corresponding RegVar scores. We calculated the optimal cutoff of RegVar scores by maximizing the sum of specificity and sensitivity. We found that a major proportion (84.5-94.0%) of the DHS-supported eVariants were correctly classified and a significant subset (24.6-39.1%) of background variants were assigned with RegVar scores above the cutoffs (**Figure 4A** and Figure S18). This result suggests that a considerable portion of variants in human genome can function as regulatory variants.

**Figure 4.**
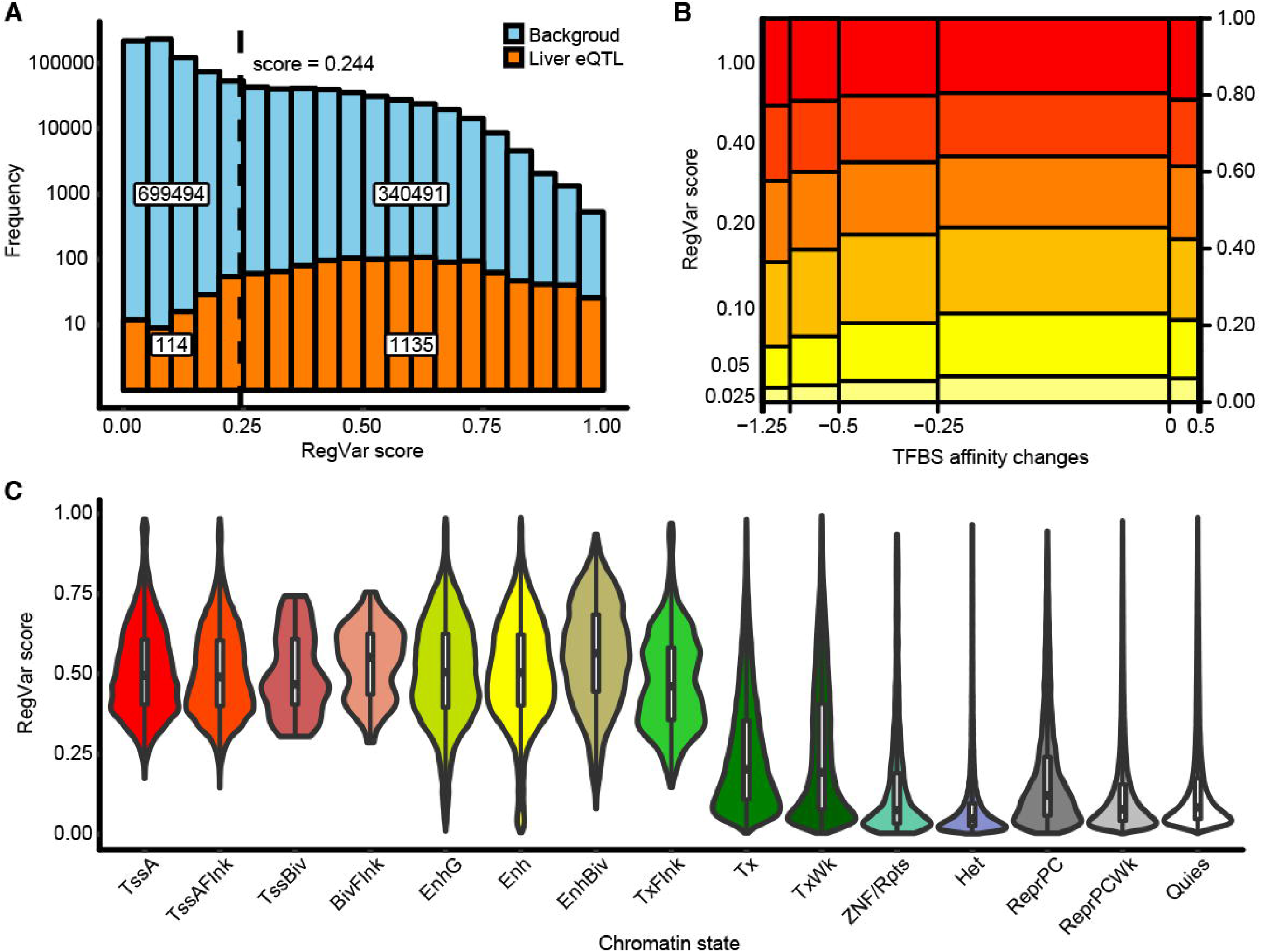
RegVar scores across all variants in chromosome 22 annotated in the integrated liver RegVar model. **A**. Histogram showing the RegVar scores distribution across all SNVs in chromosome 22 (*N* = 1,039,985) (lightblue) and SNVs in GTEx liver eQTLs (orange). Dashed line indicates the optimal cutoff score in liver training set. Numbers of variants blow or above the cutoff score are embedded. **B**. Spine plot showing the correlation between RegVar scores and SNV-caused TFBS affinity changes. **C**. Violin plots showing the RegVar score distributions across 15 chromatin states (BivFlnk, flanking bivalent TSS/enhancers; Enh, enhancers; EnhBiv, bivalent enhancers; EnhG, genic enhancers; Het, heterochromatin; ReprPC, repressed PolyComb; ReprPCWk, weak repressed PolyComb; Quies, quiescent/low; TssA, active TSS; TssAFlnk, flanking active TSS; TssBiv, bivalent/poised TSS; Tx, strong transcription; TxFlnk, transcription at gene 5’ and 3’; TxWk, weak transcription; ZNF/Rpts, ZNF genes & repeats). Embedded boxplots indicate medians (center bars), and the first and third quartiles (lower and upper hinges).

To further investigate the distribution of RegVar scores across different functional genome regions, we mapped all annotated variants across 15 chromatin states produced by ChromHMM [22] in liver and showed that variants at active/bivalent promoters and enhancers usually have higher RegVar scores, while variants at repressed and heterochromatin regions usually have lower scores (**Figure 4C**). This is expected since most variants exert their effects through alteration of key regulatory DNA elements [33]. Also, we observed a clear correlation between RegVar scores and SNV-caused loss-of-function (*ΔAffinity* <= 0, ANOVA *F* = 422.6, *P* = 0) or gain-of-function (*ΔAffinity* >= 0, ANOVA *F* = 23.62, *P* = 5.52E-11) of TFBSs (**Figure 4B**), which means the extents of TFBS affinity alteration are positively correlated with the probabilities of the causing variants to be functional.

### Tissue specificity of RegVar scores

To evaluate the tissue specificity of the predicted regulatory variants, we applied the integrated models in all 17 tissue types to randomly selected SNVs (*n* = 100,000) across the human genome. K-means clustering of the RegVar score percentiles of these SNVs identified 22 variant clusters, and one cluster was considered to be enriched in a specific tissue if it was endowed with a K-means center percentile larger than the percentile of the cutoff score in the corresponding tissue. We then identified 8, 11, and 3 clusters of non-functional, tissue-specific regulatory, and tissue-shared regulatory variants, from clusters which were enriched in 0, 1-3, and >=12 tissues (there was no cluster enriched in 4-11 tissues), respectively (**Figure 5A** and Figure S19). Using four epigenetic marks (DHS, H3K4me1, H3K27ac and H3K4me3) as hallmarks of chromatin states, we showed that two clusters of tissue-shared variants assigned with high RegVar scores (C6, C14) presented active promoter marks (DHS, H3K4me1, H3K27ac, and H3K4me3) across all tissues, indicating they were enriched at tissue-shared promoters. In contrast, the tissue-shared cluster assigned with moderate RegVar scores (C2) presented active enhancer marks (DHS, H3K4me1, and H3K27ac) across all tissues, indicating their enrichment at tissue-shared enhancers. We also found that most of the tissue-specific clusters presented active enhancer marks specifically in the corresponding tissues, indicating their enrichment at tissue-specific enhancers (**Figure 5B**). These results demonstrate the power of RegVar in measuring the tissue-specific impact of regulatory variants.

**Figure 5.**
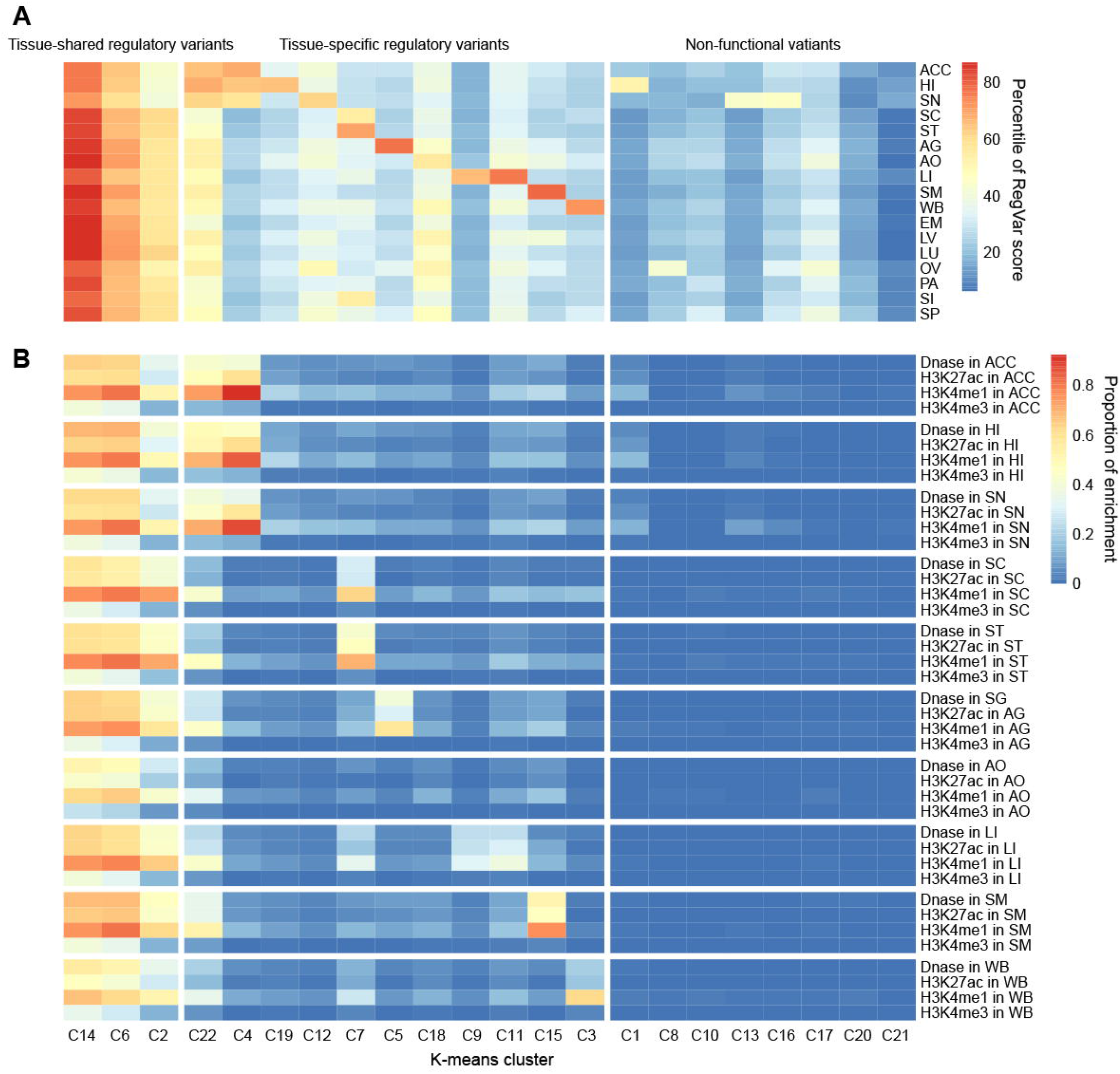
Tissue-shared and tissue-specific regulatory variants and non-functional variants identified in K-means clustering. **A**. RegVar score percentiles for different clusters of variants (*N* = 100,000) annotated with the integrated RegVar models in 17 tissues (ACC, anterior cingulate cortex; AG, adrenal gland; AO, aorta; EM, esophagus mucosa; HI, hippocampus; LI, liver; LU, lung; LV, left ventricle; OV, ovary; PA, pancreas; SC, sigmoid colon; SI, small intestine; SM, skeletal muscle; SN, substantia nigra; SP, spleen; ST, stomach; WB, whole blood). **B**. Enrichment proportion of different clusters of variants in genome regions with four epigenomic annotations (DHS, H3K4me1, H3K4me3, and H3K27ac) in 10 selected tissues.

### Prioritization of noncoding pathogenic variants in HGMD

We further extended the framework of RegVar to prioritize noncoding pathogenic variants. We used a simplified pathogenic RegVar model to learn the features of noncoding pathogenic variants collected from the Human Gene Mutation database (HGMD) [34]. We extracted disease-associated variants from the December 2016 release of HGMD public dataset. Small indels and variants overlapping any coding sequence (as annotated in RefSeq genes from the UCSC Genome Browser) or essential splice sites (as annotated in GWAVA [4]) were filtered out. After mapping remaining variants to all genomic annotations (Supplementary Methods), a final set of 2078 disease-associated variants were used as the positive set of pathogenic noncoding variants. For negative datasets, three datasets were constructed, including (i) *random-variant* set of random SNVs sampled from the whole genome; (ii) *distance-control-variant* set of random SNVs sampled from variants matched to the pathogenic ones by the exact distance-to-nearest TSS (not necessarily near the same TSSs as the pathogenic variants); (iii) *neighboring-variant* set of random SNVs located <= 1 kb from the pathogenic ones. We selected a sample ratio at 1:10 due to the small sample size of pathogenic variants, and negative variants overlapping any coding sequence or essential splice sites were further filtered out. We then constructed the pathogenic RegVar model on those different negative datasets. We found that RegVar obtained superior capability in the *random-variant* and *neighboring-variant* sets. The performances of regBase (AUC = 0.879), GWAVA (AUC = 0.874), IW-scoring (AUC = 0.871) were comparable to RegVar’s (AUC = 0.885) in the *random-variant* set, and regBase (AUC = 0.704) was comparable to RegVar (AUC = 0.707) in the *neighboring-variant* set. In the *distance-control-variant* set, regBase exhibited slight outperformance (AUC = 0.845), followed by RegVar (AUC = 0.816) and IW-scoring (AUC = 0.805) (**Figure 6**). These results demonstrated the competence of the RegVar framework in discriminating between pathogenic and benign variants.

**Figure 6.**
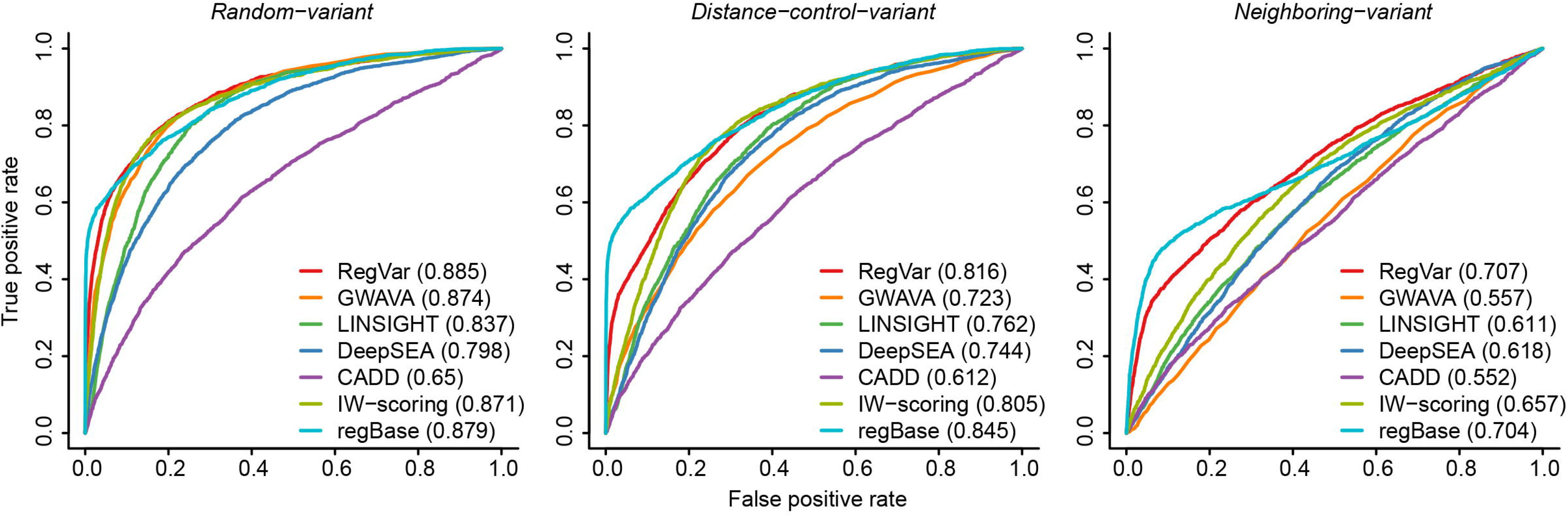
ROC curves of seven computational methods distinguishing noncoding pathogenic variants from different backgrounds. Positive samples were from the of HGMD noncoding pathogenic variants (*N* = 2078). Negative samples were from either random selected variants (*random-variant* set) (left), matched variants by the exact distance-to-nearest TSS (not necessarily near the same TSS as each pathogenic variant) (*distance-control-variant* set) (middle), or neighboring variants located adjacent (<= 1 kb) to the positive ones (*neighboring-variant* set) (right). Because GWAVA was trained on the HGMD noncoding mutations, we filtered out GWAVA training positive-variants in evaluating its performance. TIVAN and ExPecto results are not shown because they only provides tissue-specific regulatory variants prioritization scores.

We then explored the feature importance of the above three models (Supplementary Methods). We found that sequential profiles were the most important feature set in all three models, illustrating their prominent role in discriminating noncoding pathogenic variants from different backgrounds; epigenetic and evolutionary profiles were ranked second in the *random-variant* and *neighboring-variant* models, respectively (Figure S20), which demonstrated their specific facility in separating noncoding pathogenic variants from a global and local genome regions, respectively.

## Discussion

Noncoding variants play a prominent role in many diseases and complex traits through various intricate mechanisms [35, 36]. Nevertheless, variants would exert their effects by affecting the expression of specific genes. It is a great challenge to link regulatory variants, especially in long distance, to their target genes. Here we show that through jointly learning the genomic patterns of variants and genes, RegVar provides helpful information for mapping regulatory variants to their target genes. We expect RegVar can contribute to current limited understanding of genetic architecture of human genome and help to uncover novel molecular mechanisms underlying complex traits and diseases.

A number of methods have been developed for measuring the consequence and importance of noncoding variants. Though differing from each other in the underlying intuitions and specific algorithm frameworks, they mainly focused on predicting the pathogenic effect of variants. Therefore a vast number of noncoding variants with smaller regulatory effects would be neglected. Here we demonstrated the unique ability of RegVar to prioritize regulatory variants against different backgrounds. We found that in the *random-variant*, *mirrored-variant*, and *random-gene* datasets, RegVar obtained accurate and robust prediction capability; in the *neighboring-variant* dataset, RegVar exhibited relatively weak prediction power, but still superior to existing methods. These results demonstrate RegVar as an integrated model to identify genome-wide regulatory variants, and it may be not suitable for fine-mapping studies in limited regions. Applying RegVar to all SNVs in chromosome 22, we show that there is a considerable portion of variants across the wide genome showing large probabilities with which to regulate the expression of certain target genes. The reason they have not been reported may be that their effects are too subtle to be detected, coupled with limited sample sizes and low statistical power.

## Code availability

The RegVar online server is freely available at http://regvar.cbportal.org/. Downloadable datasets and source code to run RegVar on local personal computers and scripts to generate figures in the manuscript are also provided at the RegVar website.

## CRediT author statement

**Hao Lu**: Methodology, Investigation, Software, Visualization, Writing - Original Draft, Writing - Review & Editing. **Luyu Ma**: Methodology, Investigation, Visualization. **Cheng Quan**: Investigation, Software. **Lei Li**: Methodology, Investigation, Visualization. **Yiming Lu**: Conceptualization, Methodology, Investigation, Visualization, Writing - Original Draft, Writing - Review & Editing. **Gangqiao Zhou**: Conceptualization, Supervision. **Chenggang Zhang**: Conceptualization, Supervision. All authors read and approved the final manuscript.

## Competing interests

The authors have declared no competing interests.

## Acknowledgements

This work was supported by the General Program of the National Natural Science Foundation of China (Grant No. 31771397) and the Beijing Nova Program (Grant No. 20180059).

## Supplementary material

**Figure S1.**
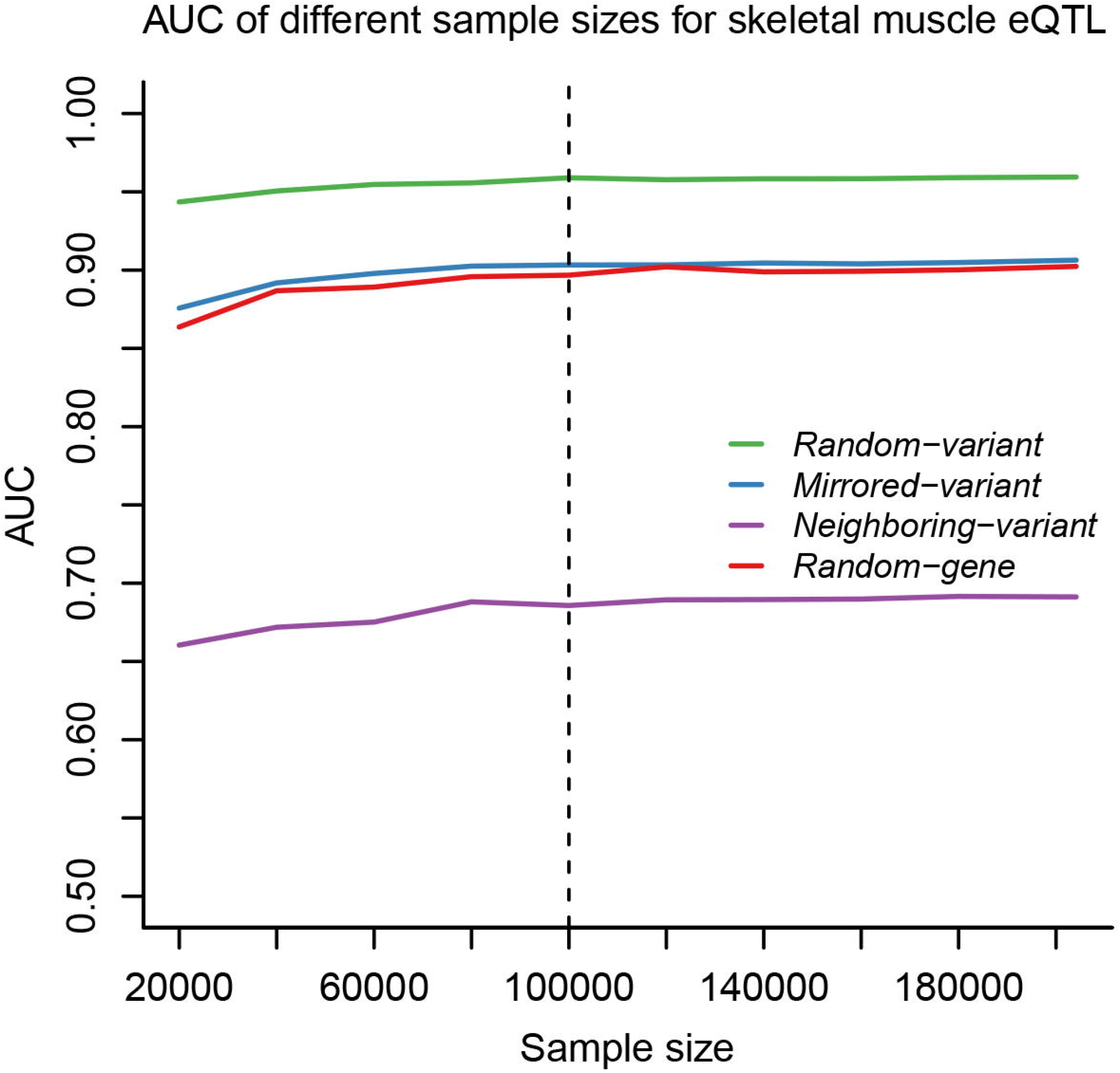
AUCs of RegVar models with different sample sizes. Positive samples were from the GTEx skeletal muscle eQTLs. Sample sizes were set from 20,000 to 180,000, step by 20,000, and the full sample size (*N* = 204,124).

**Figure S2.**
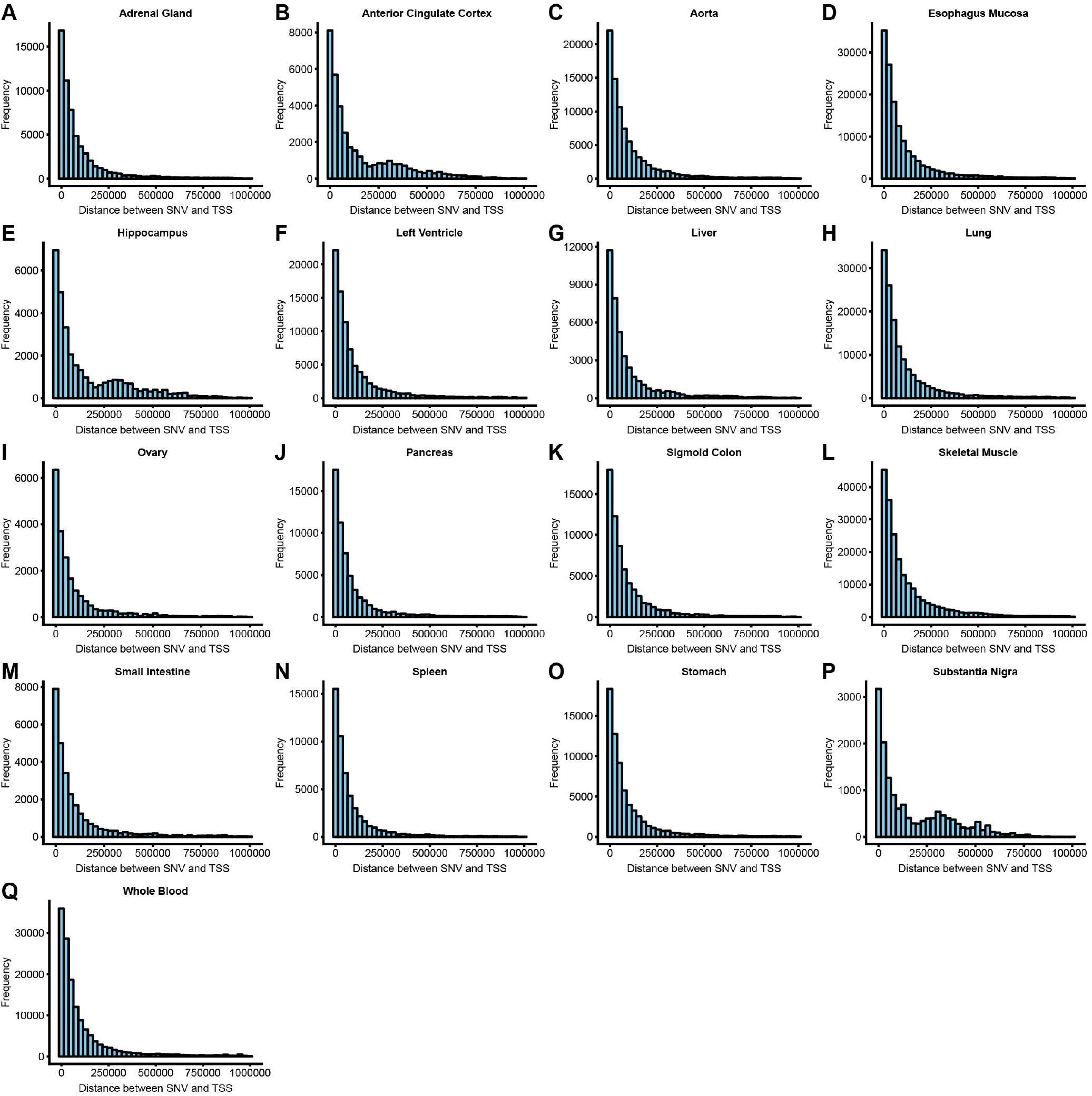
Histogram showing the distance between variant loci and TSS. Results are shown for samples from the GTEx eQTLs from 17 tissues (**A-Q**).

**Figure S3.**
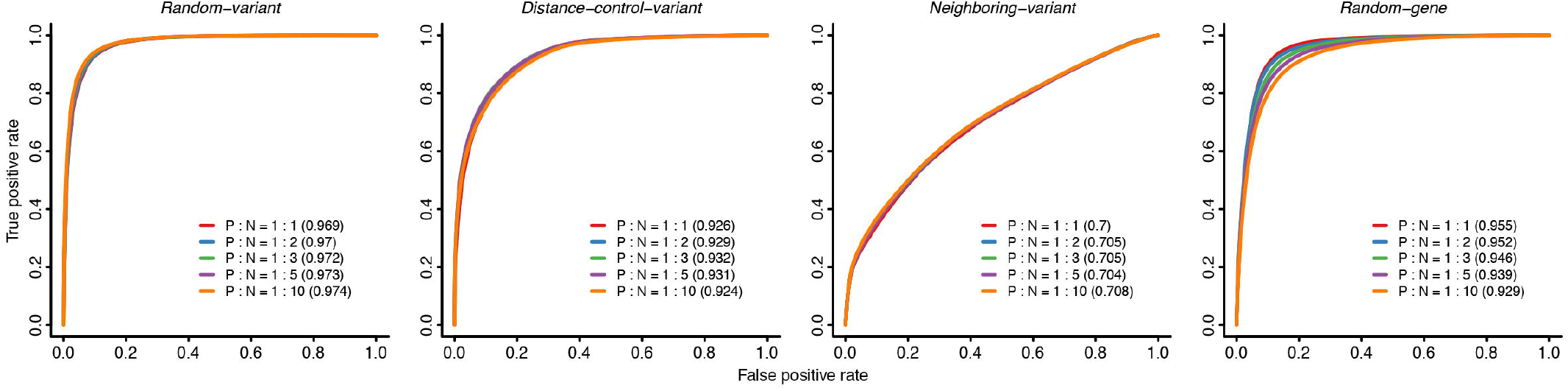
ROC curves for 10-fold cross-validation experiments at different ample ratios between positive and negative datasets. Positive samples were from the GTEx liver eQTLs. Negative samples were selected at the ratio of 1:1, 1:2, 1:3, 1:5, and 1:10 between positive (P) and negative (N) datasets.

**Figure S4.**
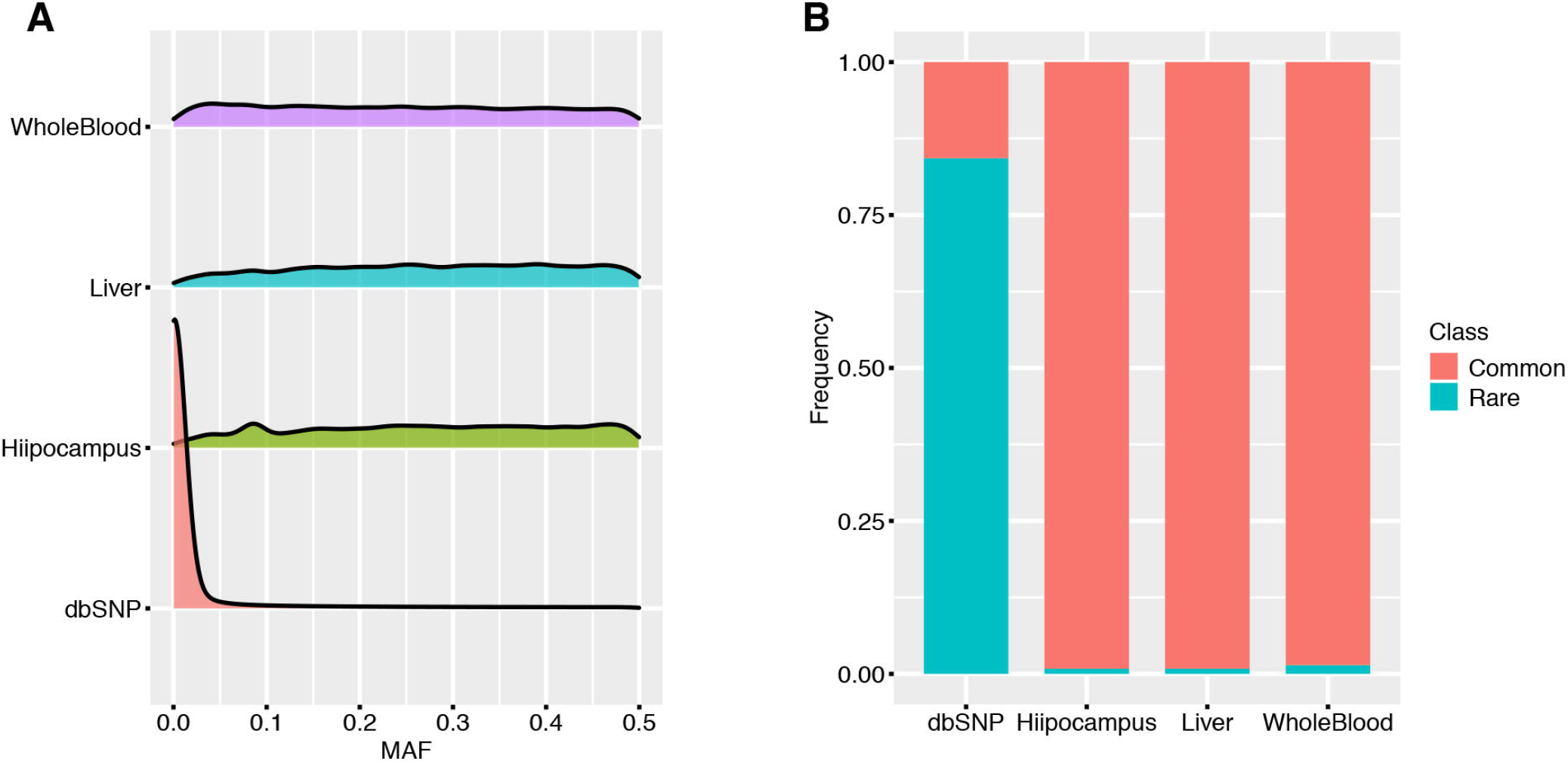
MAF distribution (A) and common and rare variant proportion (B) in dbSNP variants and GTEx eQTLs.

**Figure S5.**
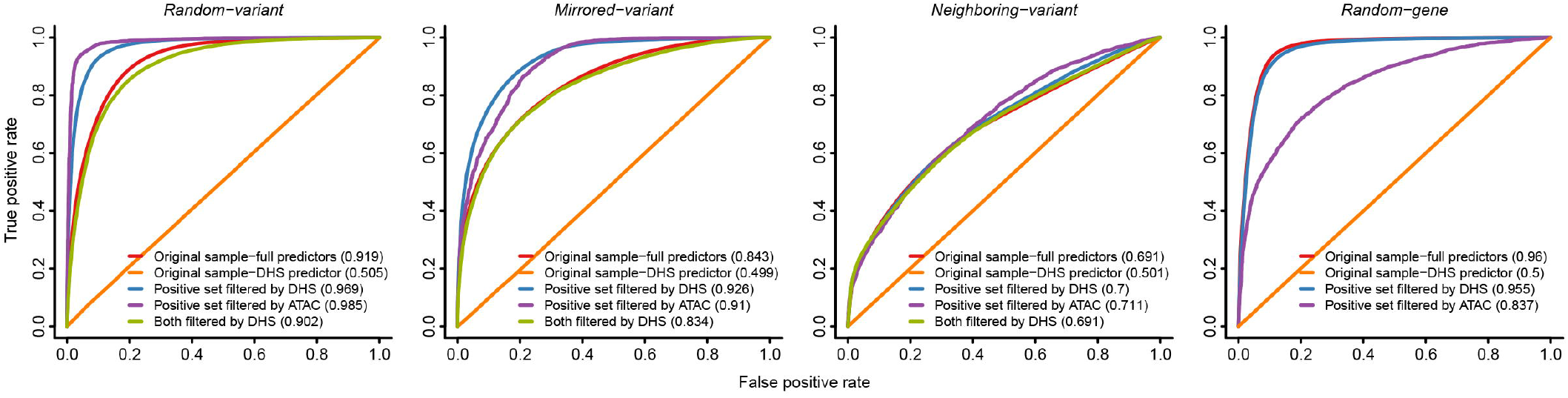
ROC curves of RegVar models trained on samples with different filters. Results are shown for models trained on the original GTEx liver eQTLs (*N* = 268,673) with all feature sets as predictors and with DHS peaks alone as a predictor, RegVar models trained on the GTEx liver eQTLs filtered by DHS peaks (*N* = 41,636) and filtered by ATAC profiles (*N* = 3731), and RegVar models trained on both the GTEx liver eQTLs and controls filtered by DHS peaks.

**Figure S6.**
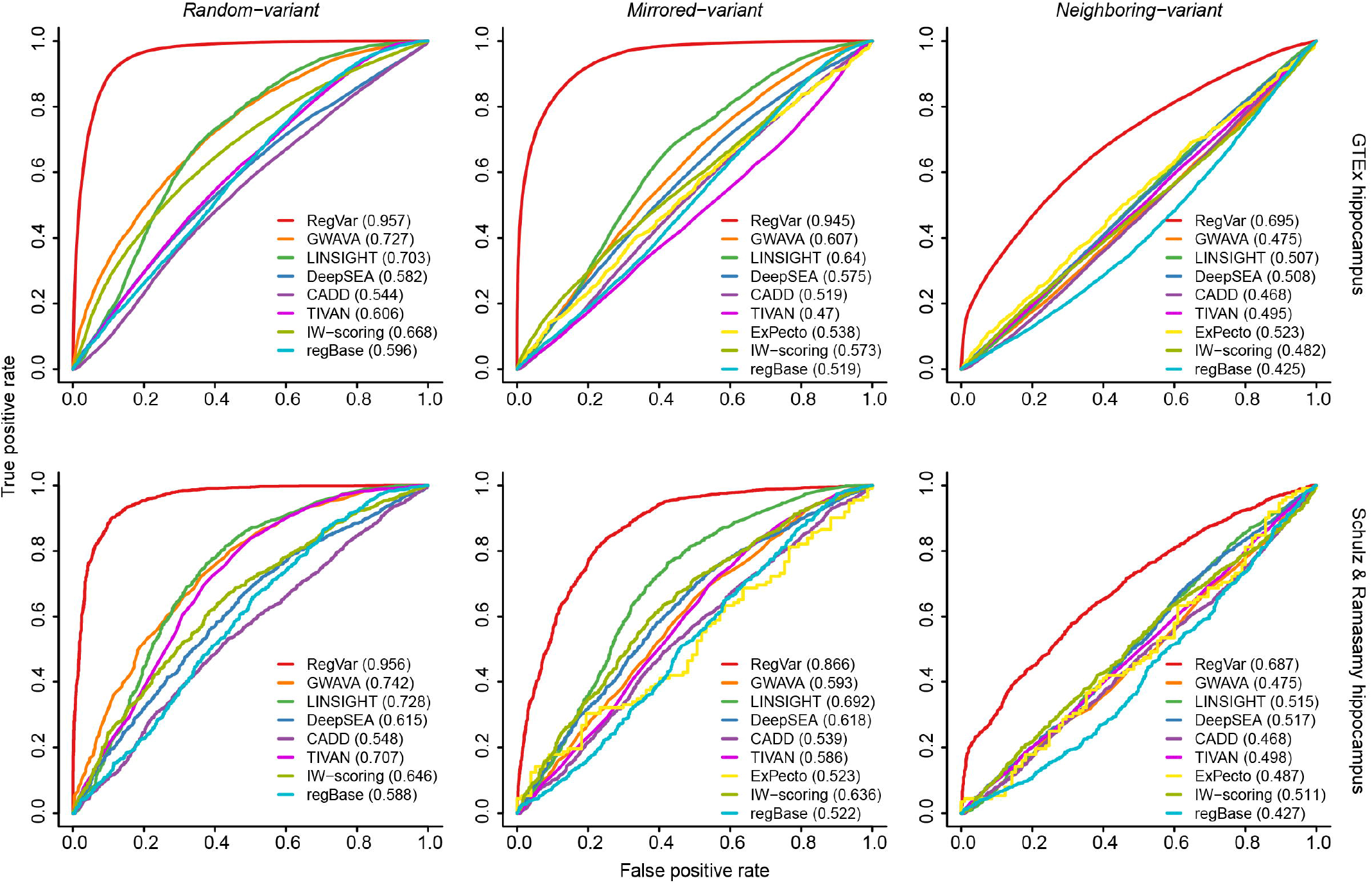
ROC curves of nine computational methods distinguishing regulatory variants from different backgrounds in hippocampus. Results are shown for ROC curves from 10-fold cross-validation experiments in the GTEx hippocampus eQTL dataset (top) and from external evaluation experiments in Schulz and Ramasamy hippocampus eQTL dataset (bottom). Plots are similar to Figure 3.

**Figure S7.**
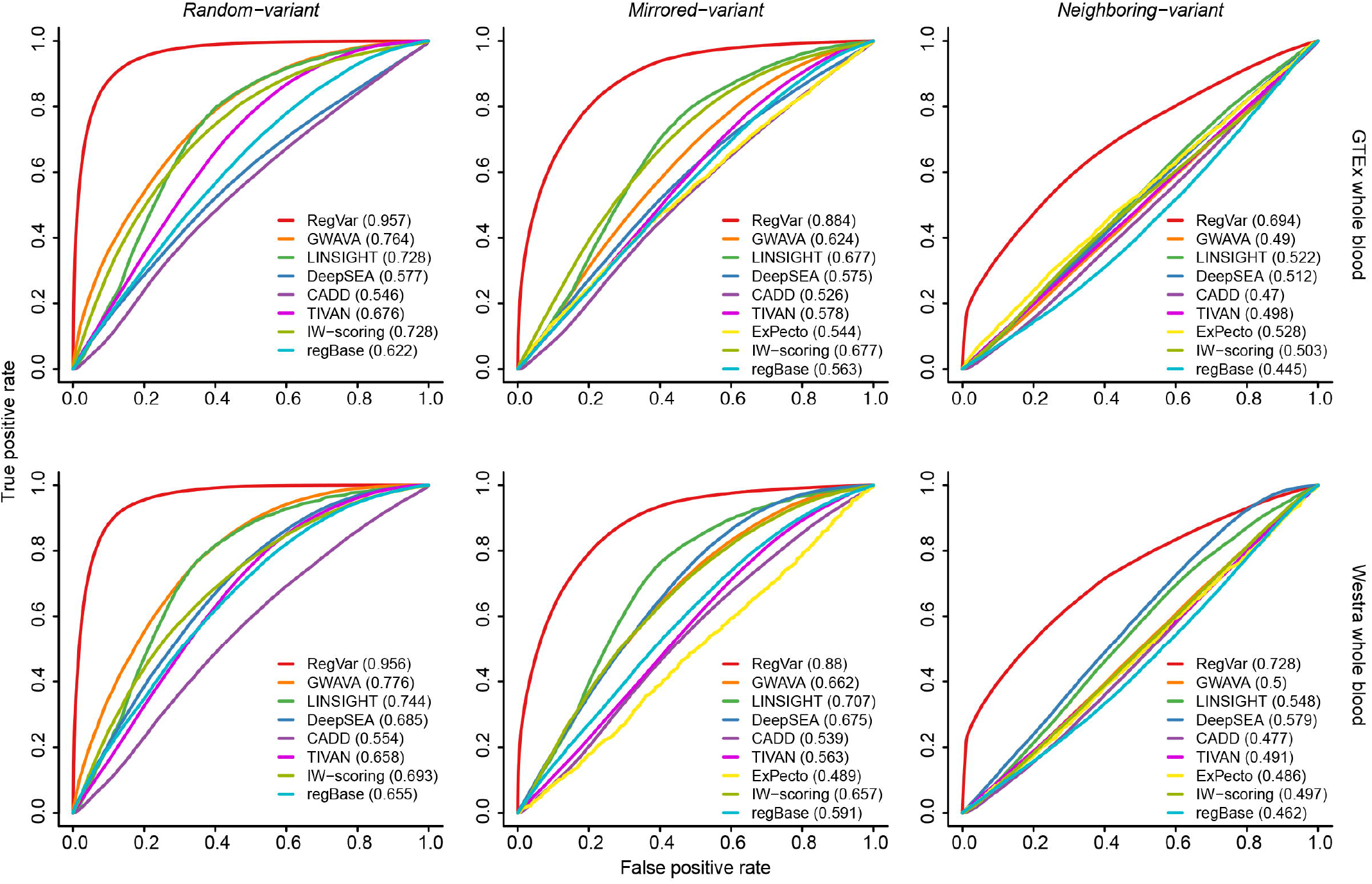
ROC curves of nine computational methods distinguishing regulatory variants from different backgrounds in whole blood. Results are shown for ROC curves from 10-fold cross-validation experiments in the GTEx whole blood eQTL dataset (top) and from external evaluation experiments in Westra whole blood eQTL dataset (bottom). Plots are similar to Figure 3.

**Figure S8.**
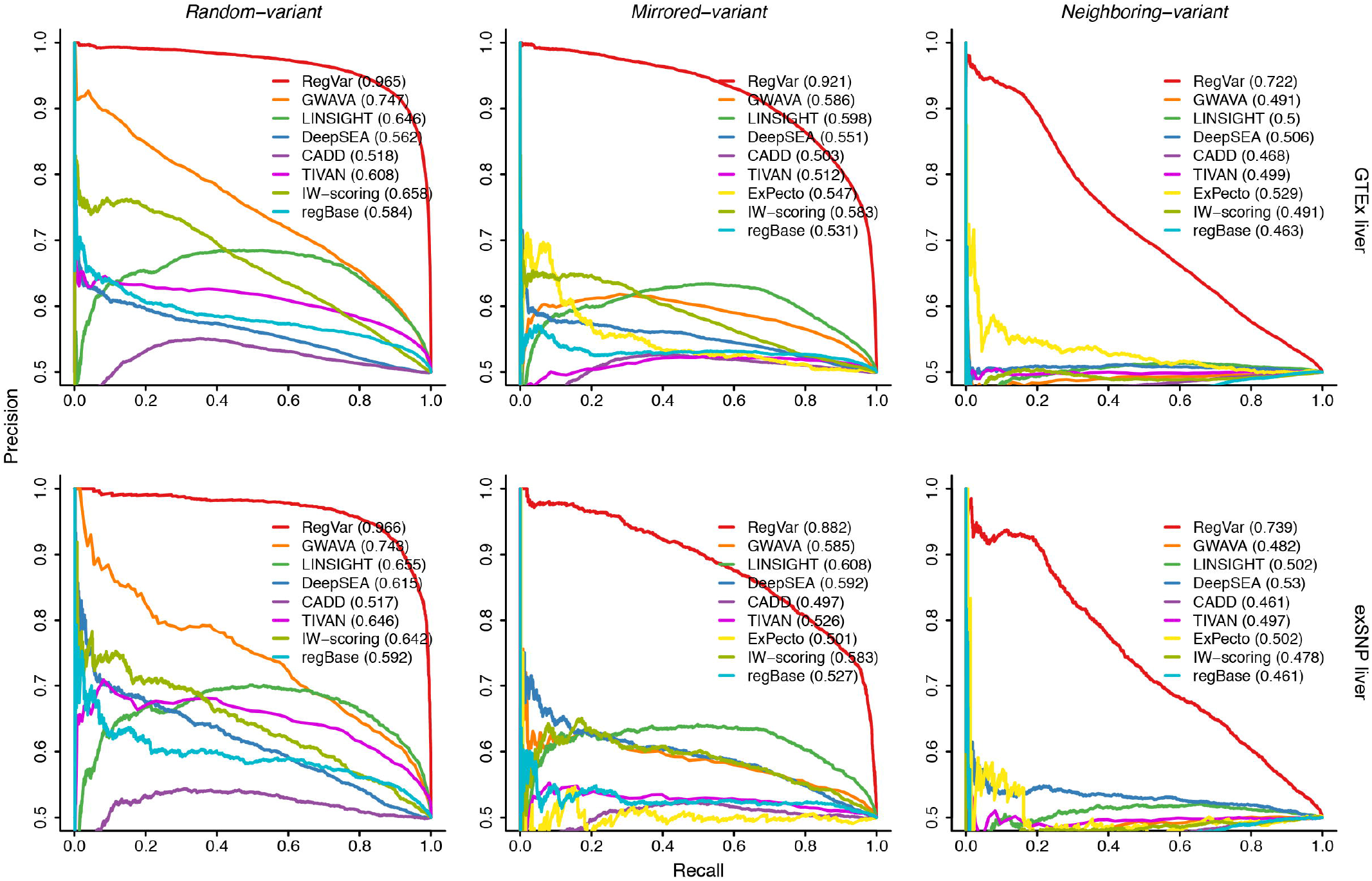
PRC curves of nine computational methods distinguishing regulatory variants from different backgrounds in liver. Results are shown for PRC curves for the same result as Figure 3.

**Figure S9.**
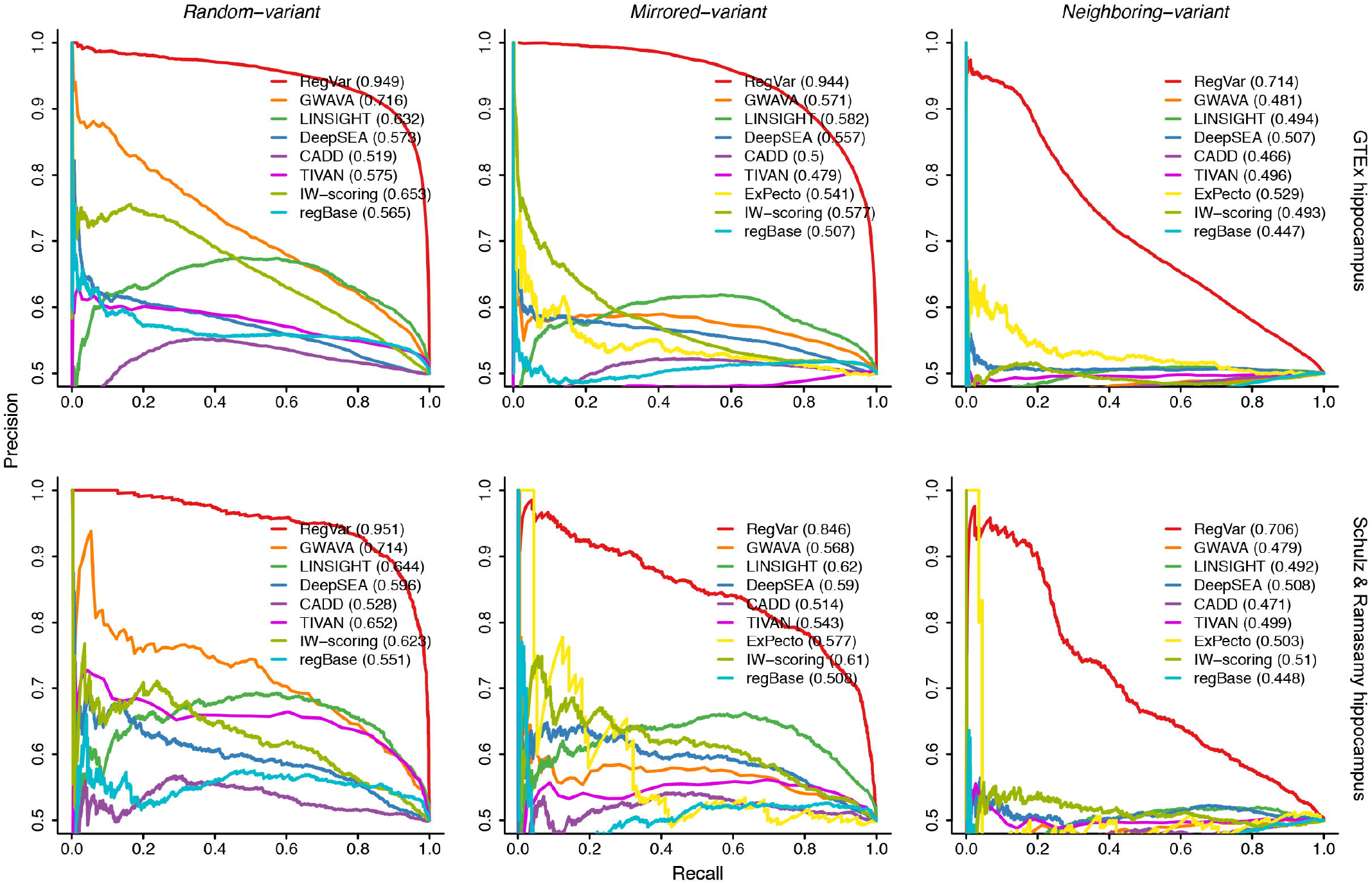
PRC curves of nine computational methods distinguishing regulatory variants from different backgrounds in hippocampus. Results are shown for PRC curves for the same result as Supplementary Figure S6

**Figure S10.**
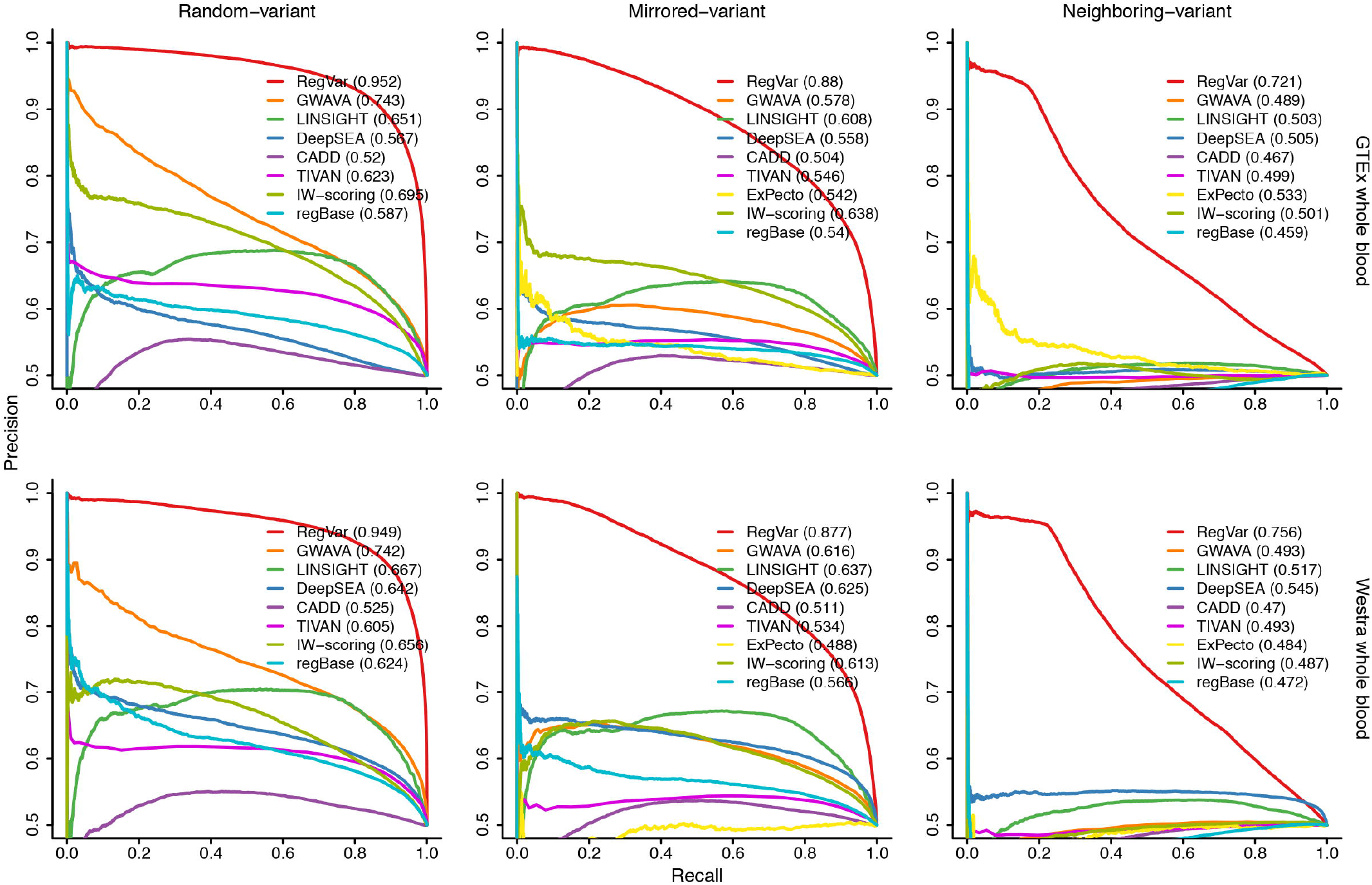
PRC curves of nine computational methods distinguishing regulatory variants from different backgrounds in whole blood. Results are shown for PRC curves for the same result as Supplementary Figure S7

**Figure S11.**
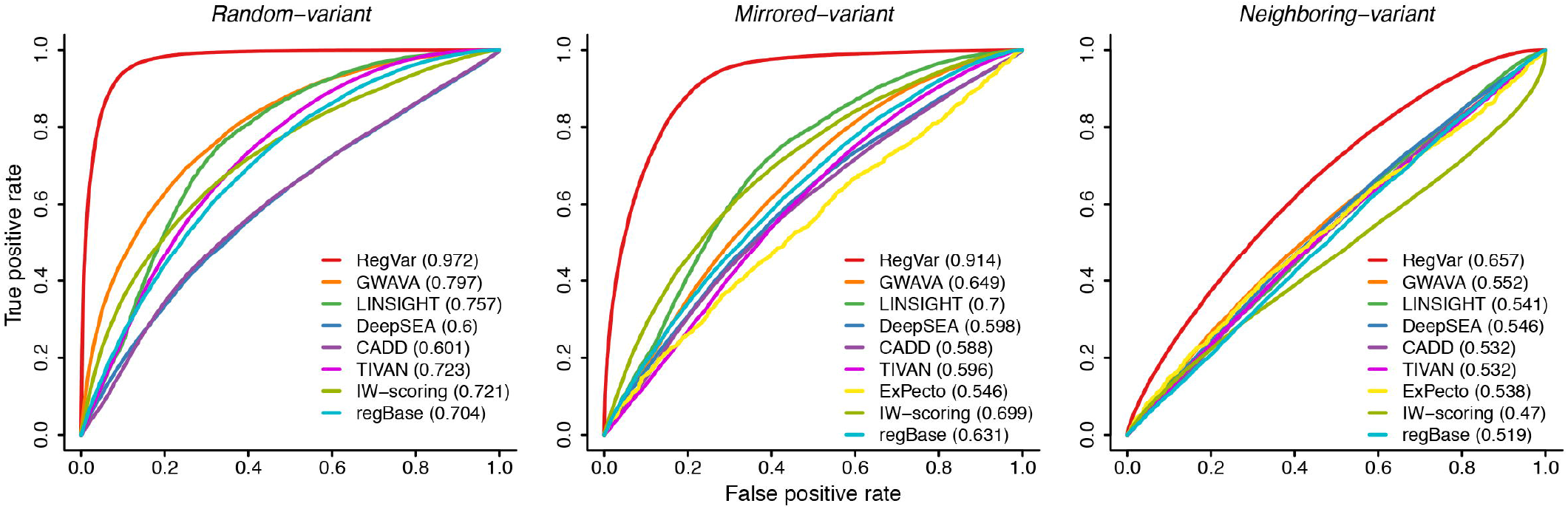
ROC curves of nine computational methods distinguishing regulatory variants in liver from MAF-matched variants. Positive samples were from the GTEx liver eQTLs (*N* = 41,636). Negative samples in three control datasets were randomly selected from MAF-matched variants to the positive ones. Plots are similar to Figure 3.

**Figure S12.**
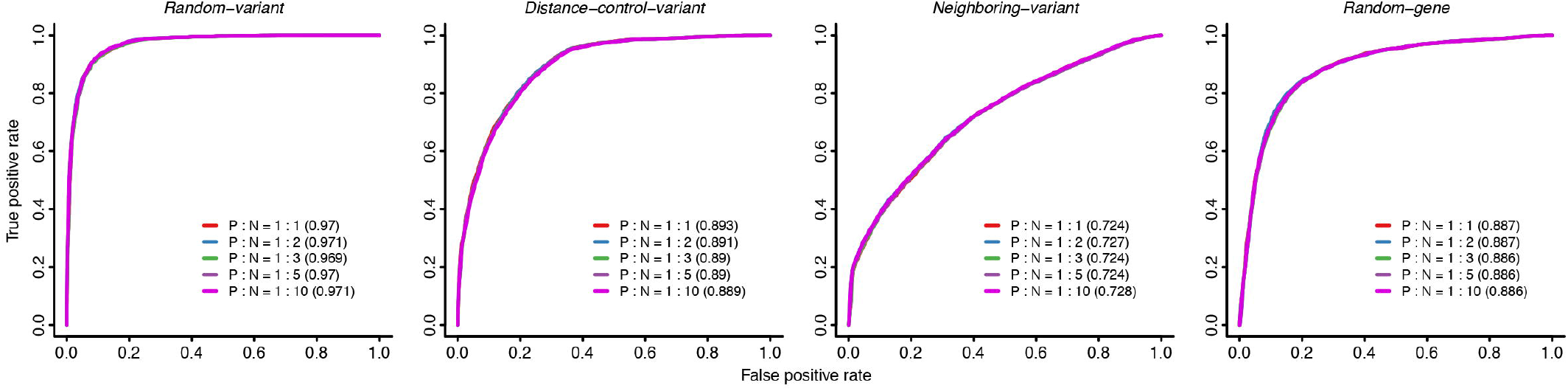
ROC curves of RegVar distinguishing regulatory variants from different backgrounds in liver at different sample ratio. Positive samples were from the exSNP liver eQTL dataset. Negative samples were selected at the ratio of 1:1, 1:2, 1:3, 1:5, and 1:10 between positive (P) and negative (N) datasets.

**Figure S13.**
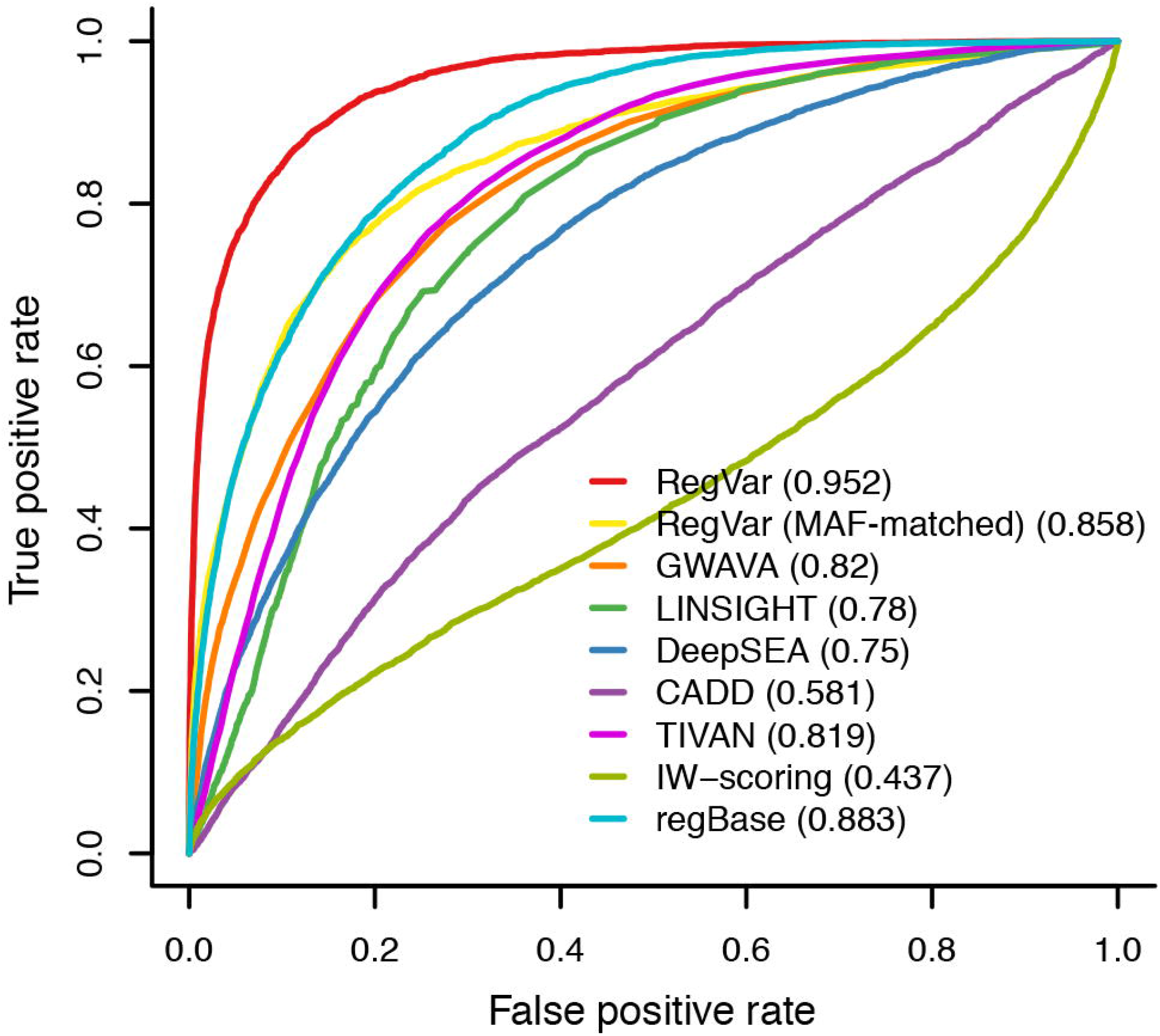
ROC curves of nine computational methods distinguishing regulatory variants in Brown liver eQTLs. Positive and negative samples were from the Brown liver eQTLs complied by regBase.

**Figure S14.**
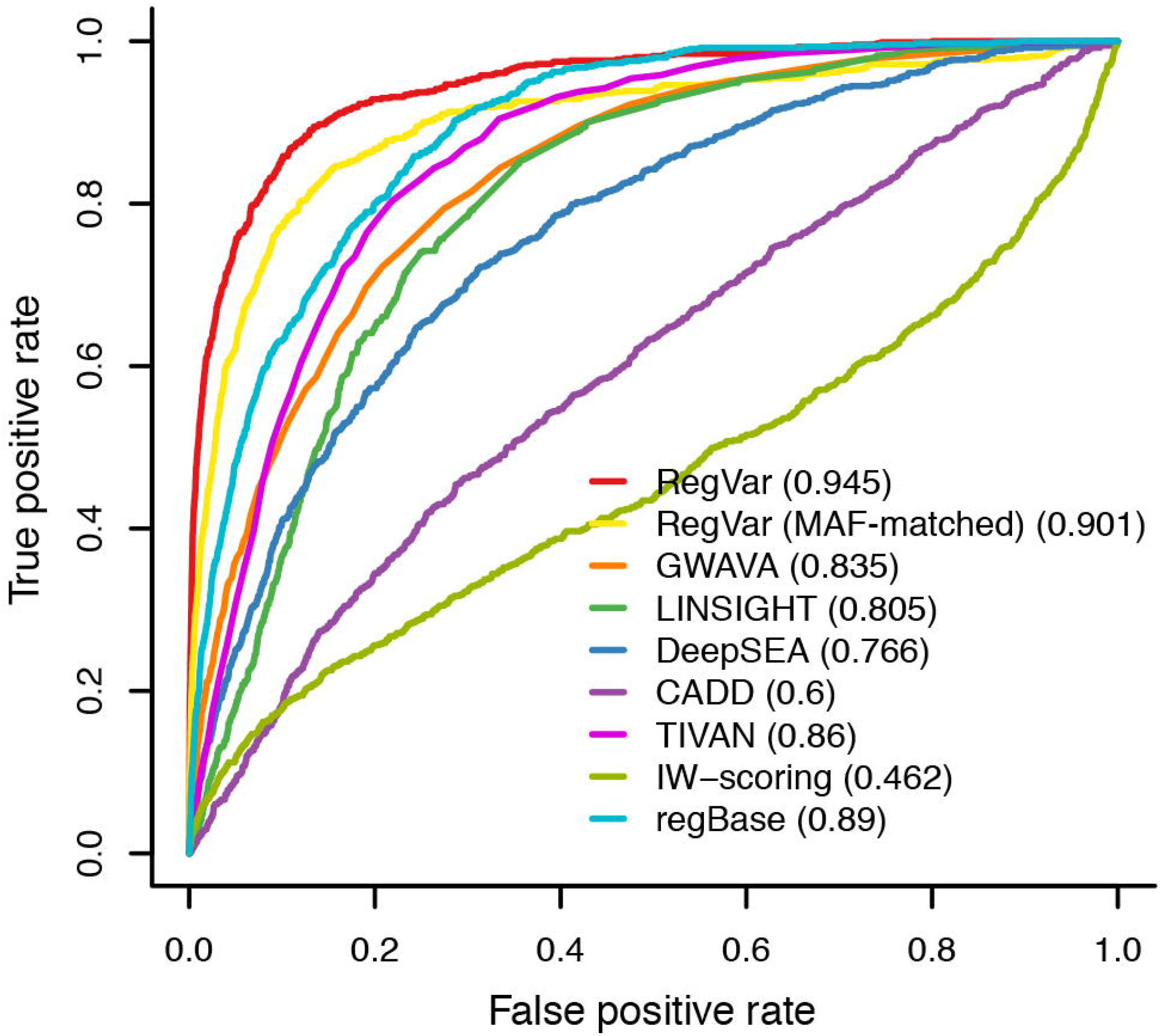
ROC curves of nine computational methods distinguishing regulatory variants in Brown blood eQTLs. Positive and negative samples were from the Brown blood eQTLs complied by regBase.

**Figure S15.**
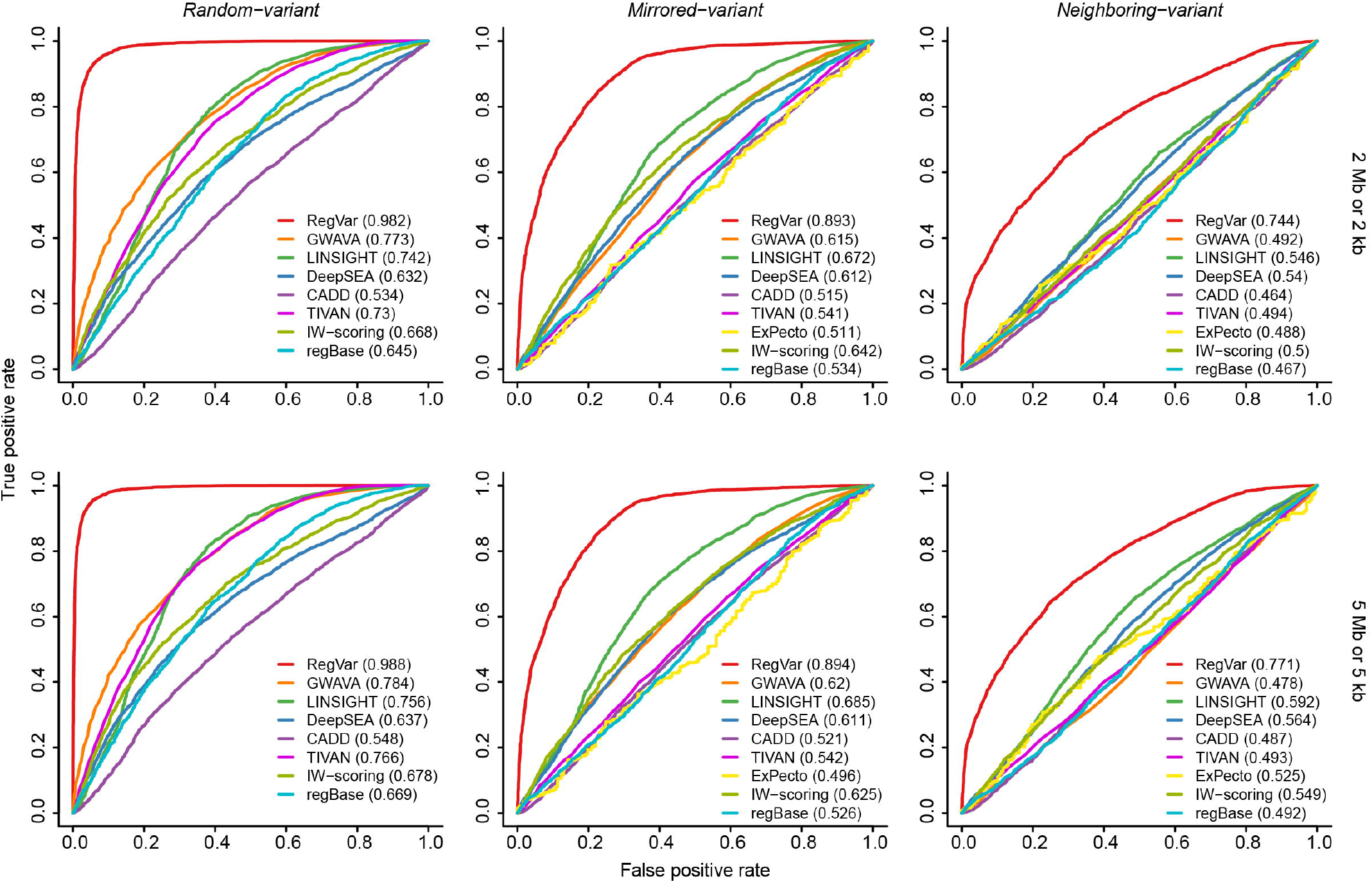
ROC curves of nine computational methods distinguishing regulatory variants in the exSNP liver eQTLs from different backgrounds. Positive samples were from the exSNP liver eQTL dataset (*N* = 4307). Negative samples in three control datasets were randomly selected from wider genome regions: (i) *random-variant* set comprising of random SNVs located <= 2 (Top) and 5 (bottom) Mb from the eGene TSS; (ii) *mirrored-variant* set comprising of random SNVs with a distance error <= 2 (Top) and 5 (bottom) kb; (iii) *neighboring-variant* set comprising of random SNVs located <= 2 (Top) and 5 (bottom) kb to the positive ones. Result for *random-gene* set comprising of random gene TSSs located <= 2 and 5 Mb of the eVariants was not shown since other existing methods didn’t give prediction on potentially affected genes.

**Figure S16.**
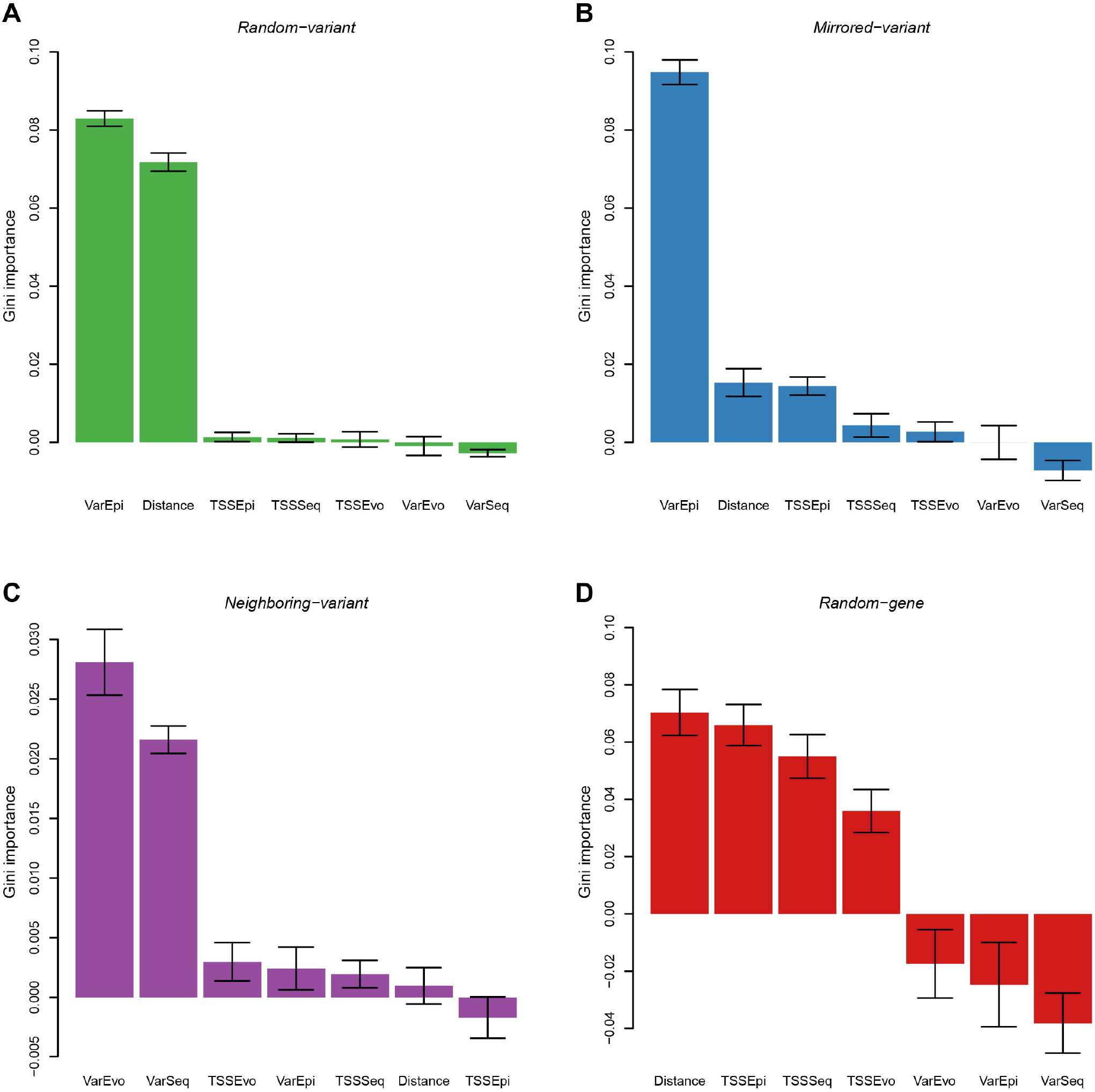
Barplots showing the relative Gini importance for different models. Results are shown for models trained on *random-variant* (**A**), *mirrored-variant* (**B**), *neighboring-variant* (**C**), and *random-gene* (**D**) datasets of the GTEx liver eQTLs (*N* = 41,636). All features were divided into seven groups: distance between variant and TSS (Distance), sequential profiles of variant (VarSeq) and TSS (TSSSeq), epigenetic profiles of variant (VarEpi) and TSS (TSSEpi), evolutionary profiles of variant (VarEvo) and TSS (TSSEvo). Error bars represents the standard error averaged over 10 times.

**Figure S17.**
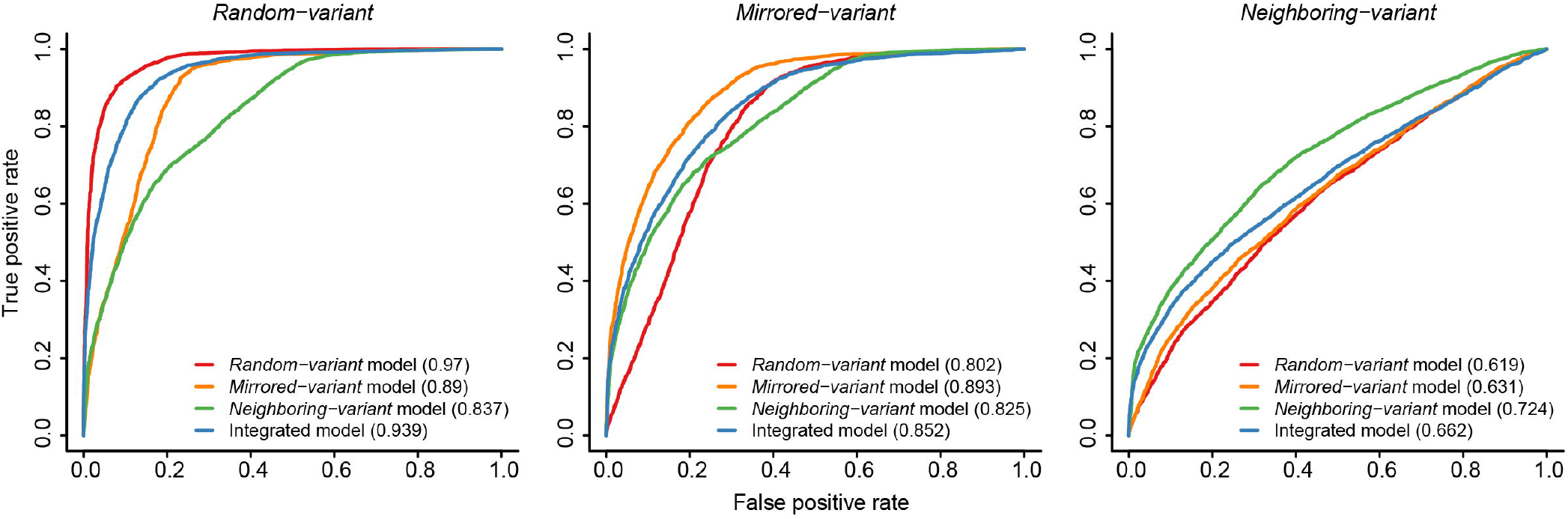
ROC curves of different RegVar models distinguishing regulatory variants in the exSNP liver eQTLs from each of the negative datasets. Positive samples were from the exSNP liver eQTLs (*N* = 4307). Models based on the *random-gene* dataset was not evaluated in this analysis, because it aimed to identify potential eGenes for each eVariant and wasn’t suitable for prioritization of positive variants.

**Figure S18.**
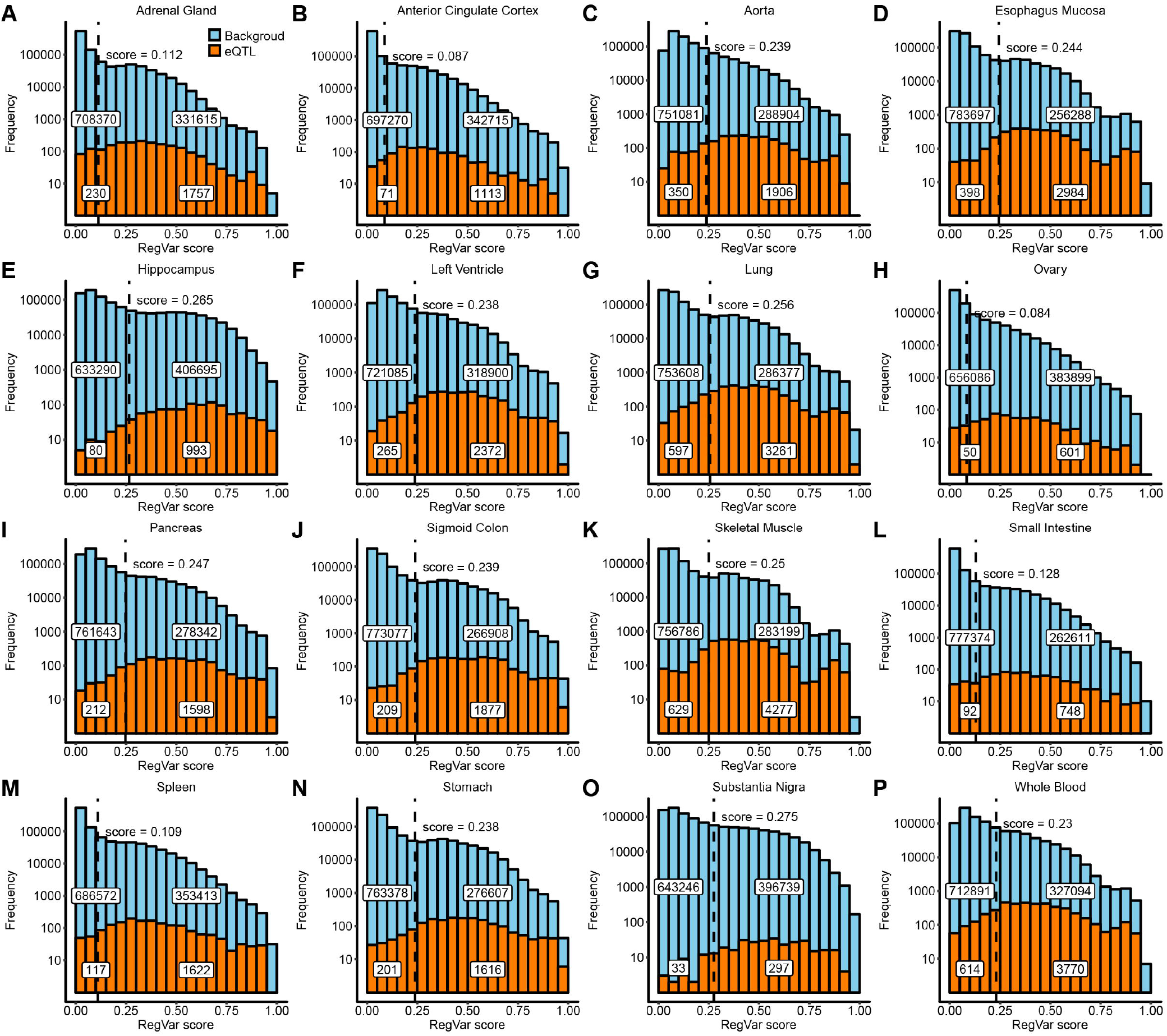
Histogram showing the RegVar scores distribution. Results are shown for all SNVs in chromosome 22 (lightblue) and SNVs in the GTEx eQTLs (orange). Variants were annotated in integrated RegVar models in 16 tissues (**A-P**). Dashed line indicates the optimal cutoff scores in the corresponding training set. Numbers of variants blow or above the cutoff scores are embedded.

**Figure S19.**
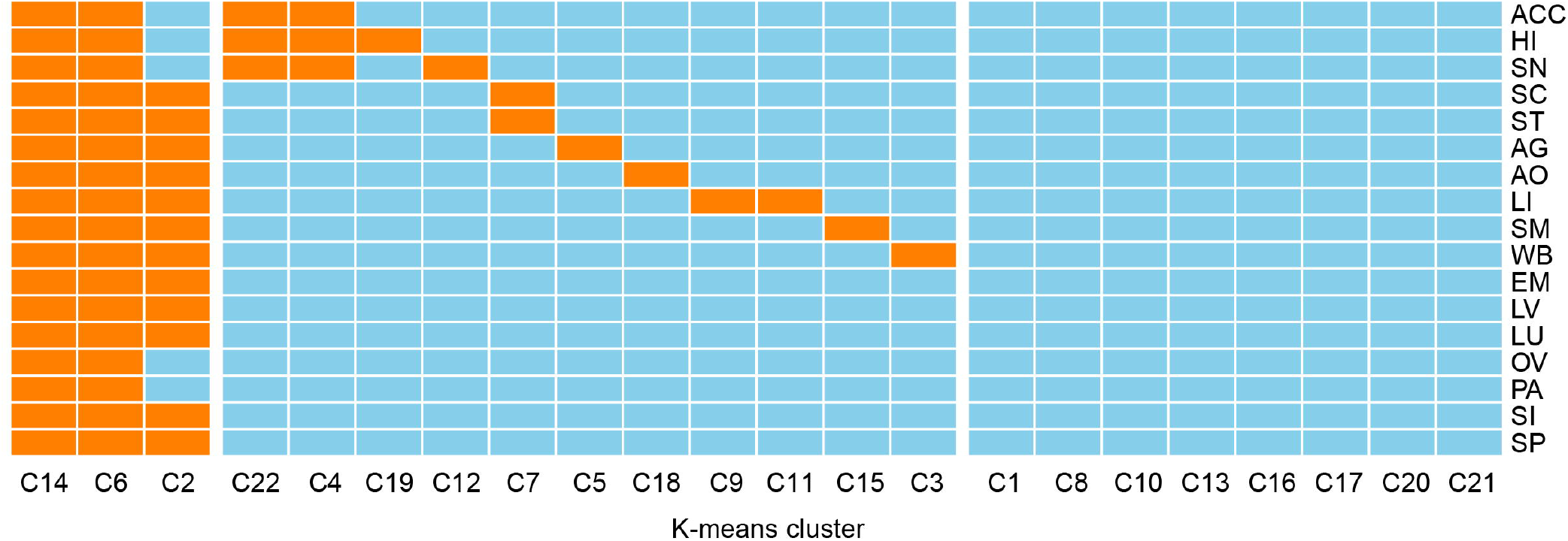
Identification of tissue-shared/tissue-specific cluster of variants. An orange cell indicate the cluster was endowed with an K-means center percentile larger than the percentile of the RegVar cutoff score in the corresponding tissue, and an skyblue cell if not. ACC, anterior cingulate cortex; AG, adrenal gland; AO, aorta; EM, esophagus mucosa; HI, hippocampus; LI, liver; LU, lung; LV, left ventricle; OV, ovary; PA, pancreas; SC, sigmoid colon; SI, small intestine; SM, skeletal muscle; SN, substantia nigra; SP, spleen; ST, stomach; WB, whole blood.

**Figure S20.**
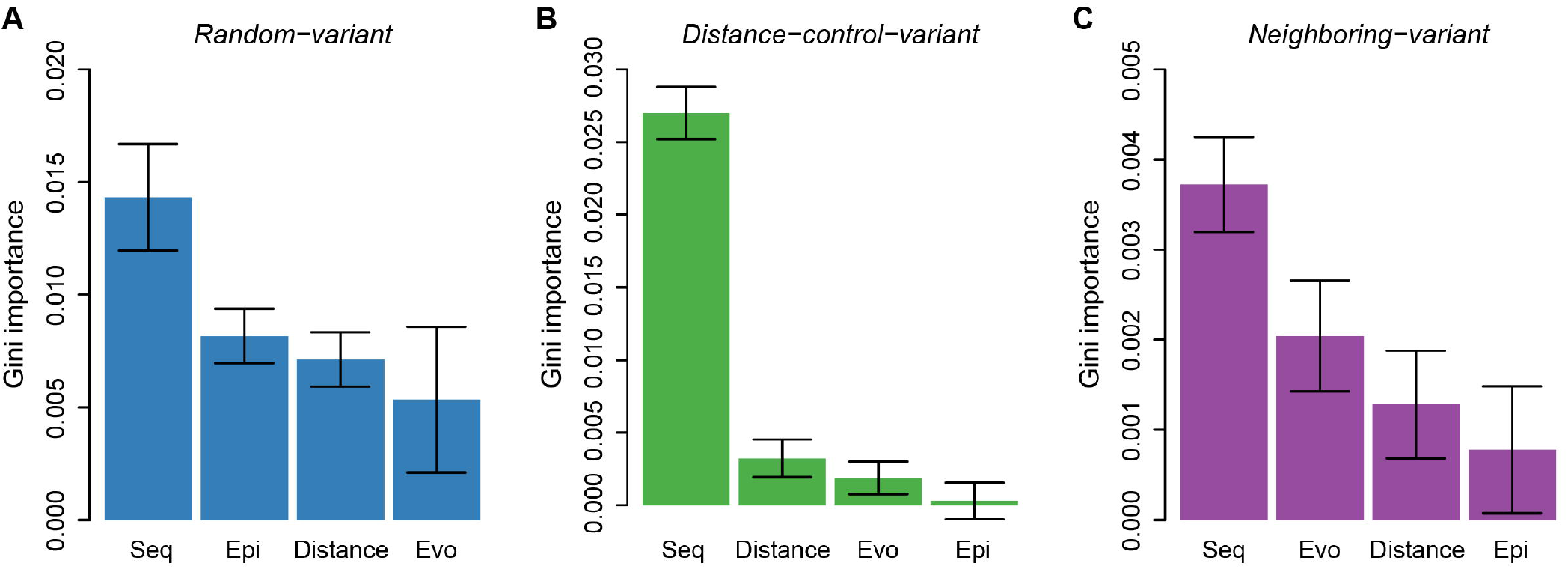
Barplots showing the relative Gini importance for different pathogenic RegVar models. Results are shown for models trained on *random-variant* (**A**), *distance-control-variant* (**B**), and *neighboring-variant* (**C**) datasets of HGMD noncoding pathogenic variants (*N* = 2,078). All features were divided into four groups: distance to the nearest TSS (Distance), and sequential (Seq), epigenetic (Epi), and evolutionary (Evo) profiles of the variant. Error bars represents the standard error averaged over 10 times.

**Table S1 Sample sizes and AUCs of the 4 training sets in 17 tissues**

**Table S2 Summary of genomic features used by RegVar**

**Table S3 AUCs under different learning rates and dropout proportions in the liver *random-variant* set**

**Table S4 AUCs under different learning rates and dropout proportions in the liver *mirrored-variant* set**

**Table S5 AUCs under different learning rates and dropout proportions in the liver *neighboring-variant* set**

**Table S6 AUCs under different learning rates and dropout proportions in the liver *random-gene* set**

**Table S7 Testing sets used for evaluation of RegVar and other methods**

**File S1 Supplementary methods**

